# Nuclear SUN1 Stabilizes Endothelial Cell Junctions via Microtubules to Regulate Blood Vessel Formation

**DOI:** 10.1101/2021.08.11.455980

**Authors:** Danielle B Buglak, Molly R Kulikauskas, Ziqing Liu, Ariel L Gold, Allison P Marvin, Andrew Burciu, Natalie T Tanke, Shea N Ricketts, Karina Kinghorn, Morgan Oatley, Bryan N Johnson, Pauline Bougaran, Celia E Shiau, Stephen L Rogers, Victoria L Bautch

**Author notes:** Corresponding author: Victoria L Bautch, PhD, Professor of Biology, Department of Biology, CB#3280, University of North Carolina at Chapel Hill, Chapel Hill, NC 27599 USA.

## Abstract

Endothelial cells line all blood vessels, where they coordinate blood vessel formation and the blood-tissue barrier via regulation of cell-cell junctions. The nucleus also regulates endothelial cell behaviors, but it is unclear how the nucleus contributes to endothelial cell activities at the cell periphery. Here we show that the nuclear-localized LINC complex protein SUN1 regulates vascular sprouting and barrier function via effects on endothelial cell-cell junction morphology and function. Loss of murine endothelial *Sun1* impaired blood vessel formation and destabilized junctions, angiogenic sprouts formed but retracted in SUN1-depleted sprouts, and zebrafish vessels lacking Sun1b had aberrant junctions and defective cell-cell connections. At the cellular level, SUN1 stabilized endothelial cell-cell junctions, promoted barrier function, and regulated contractility. Mechanistically, SUN1 depletion altered cell behaviors via the cytoskeleton without changing transcriptional profiles. Reduced peripheral microtubule density, fewer junction contacts and increased catastrophes accompanied SUN1 loss, and microtubule depolymerization phenocopied effects on junctions. Depletion of GEF-H1, a microtubule-regulated Rho activator, or the LINC complex protein nesprin-1 rescued defective junctions of SUN1-depleted endothelial cells. Thus, endothelial SUN1 regulates peripheral cell-cell junctions from the nucleus via LINC complex-based microtubule interactions that affect peripheral microtubule dynamics and Rho-regulated contractility, and this long-range regulation is important for proper blood vessel sprouting and barrier function.

**SUMMARY:** The nuclear membrane protein SUN1 promotes blood vessel formation and barrier function by stabilizing endothelial cell-cell junctions. Communication between SUN1 and endothelial cell junctions relies upon proper microtubule dynamics and Rho signaling far from the nucleus, revealing long-range cellular communication from the nucleus to the cell periphery that is important for vascular development and function.

## INTRODUCTION

Blood vessels form and expand via sprouting angiogenesis, a dynamic process whereby endothelial cells migrate from pre-existing vessels to form new conduits (Carmeliet & Jain, 2011; Wacker & Gerhardt, 2011; Kushner & Bautch, 2013; Bautch & Caron, 2015). During angiogenesis, endothelial cell-cell junctions destabilize and rearrange to allow for repolarization and migration towards pro-angiogenic cues (Esser et al., 1998; Dejana, 2004; Blum et al., 2008). Specifically, the endothelial cell adherens junction protein VE-cadherin is required for vascular sprouting and viability (Carmeliet et al., 1999; Montero-Balaguer et al., 2009; Sauteur et al., 2014; Szymborska & Gerhardt, 2018). As vessels mature, endothelial cell junctions stabilize and form a functional barrier that regulates egress of fluid and oxygen; barrier dysfunction leads to increased permeability and severe disease (Claesson-Welsh, 2015; Rho et al., 2017; Claesson-Welsh et al., 2021). Thus, regulation of endothelial cell junction stability is important developmentally and for vascular homeostasis.

Adherens junctions are key to integrating and regulating both external and internal cellular inputs from multiple sources, including the microtubule and actin cytoskeletons (Ligon et al., 2001; Shaw et al., 2007; Bellett et al., 2009; Dejana & Vestweber, 2013; Abu Taha & Schnittler, 2014). For example, increased actomyosin contractility destabilizes endothelial cell adherens junctions, and disorganized junctional actin accompanies VE-cadherin loss (Huveneers et al., 2012; Sauteur et al., 2014; Angulo-Urarte et al., 2018). VE-cadherin loss also changes microtubule dynamics, and disruption of microtubule dynamics destabilizes junctions and barrier function (Komarova et al., 2012). Coordination of inputs from the actin and microtubule cytoskeletons regulates endothelial cell barrier integrity and sprouting dynamics via small GTPases (Birukova et al., 2006; Mavria et al., 2006; Sehrawat et al., 2008, 2011; Wimmer et al., 2012; Szymborska & Gerhardt, 2018). In particular, RhoA signaling is regulated by microtubules via GEF-H1, a RhoGEF that is inactive while bound to microtubules and activated upon release, leading to RhoA activation, increased actomyosin contractility, and changes to endothelial cell barrier function (Krendel et al., 2002; Birukova et al., 2006; Birkenfeld et al., 2008). However, how the endothelial cell nucleus affects these processes is poorly understood.

The nucleus is usually found far from the cell periphery and junctions, yet it is important for functions critical to angiogenesis and vascular remodeling, such as polarity, migration, and mechanotransduction (Tkachenko et al., 2013; Guilluy et al., 2014; Graham et al., 2018), and perturbations of some nuclear membrane proteins affect transcriptional profiles (Li et al., 2017; May & Carroll, 2018; Carley et al., 2021). The <u>li</u>nker of the <u>n</u>ucleoskeleton and <u>c</u>ytoskeleton (LINC) complex is comprised of both SUN (<u>S</u>ad1p, <u>UN</u>C-84) and KASH (<u>K</u>larsicht, <u>A</u>NC-1, <u>S</u>yne/Nesprin <u>H</u>omology) proteins (Starr & Fridolfsson, 2010) that function as a bridge between the nucleus and the cytoskeleton, and also link through subnuclear lamin filaments to chromatin (Haque et al., 2006). SUN proteins localize to the inner nuclear membrane and bind KASH proteins, or nesprins, from their C-terminus and lamins at their N-terminus, thus providing a structural link from the nuclear cortex to the cellular cytoskeleton (Padmakumar et al., 2005; Haque et al., 2006; McGee et al., 2006; Stewart-Hutchinson et al., 2008). Nesprins are long spectrin-rich proteins localized to the outer nuclear envelope that bind SUN proteins via their C-terminus while N-terminally interacting indirectly with microtubules (via various motor proteins such as dynein and kinesin) and intermediate filaments (via plectins), and directly with actin via calponin homology domains (Ketema et al., 2007; Meyerzon et al., 2009; Zhang et al., 2009; Fridolfsson et al., 2010; Starr & Fridolfsson, 2010). Two mammalian SUN proteins are ubiquitously expressed, and based on functional consequences of SUN manipulations it has been posited that SUN1 regulates microtubule-based functions while SUN2 coordinates actin regulation (Zhu et al., 2017). However, *in vitro* binding studies do not reveal a SUN-nesprin specificity to account for this bias (Stewart-Hutchinson et al., 2008; Ostlund et al., 2009), so how complexes are assembled and sorted in cells is unclear. It is well-established that the LINC complex integrates external inputs sensed by focal adhesions, such as substrate stiffness, to regulate transcription (Carley et al., 2022), but how the LINC complex relays signals from the nucleus to the cell periphery is less understood.

The LINC complex functions in cultured endothelial cells, as knockdown of nesprin-3 leads to impaired endothelial polarity under flow (Morgan et al., 2011), while nesprin-2 and lamin A regulate proliferation and apoptosis in endothelial cells exposed to shear stress (Han et al., 2015). Depletion of nesprin-1 alters tension on the nucleus (Chancellor et al., 2010; Anno et al., 2012), and knockdown of nesprin-1 or nesprin-2 leads to reduced collective endothelial migration (Chancellor et al., 2010; King et al., 2014). Recent work showed compromised matrix adhesion and barrier function of cultured endothelial cells using a dominant negative KASH (Denis et al., 2021). However, less is understood about the roles of the SUN proteins in endothelial cell function.

The LINC complex is required for viability *in vivo*. Loss of both mammalian *Sun* genes is embryonic lethal due to impaired neuronal nuclear migration required for proper neuronal differentiation (Lei et al., 2009; Zhang et al., 2009). Global *Sun2* loss affects epidermal nuclear positioning and cell adhesion, leading to alopecia (Stewart et al., 2015), while global loss of both *Sun* genes impairs epidermal differentiation due to altered integrin signaling (Carley et al., 2021). A role for the LINC complex in mechanotransduction *in vivo* is suggested by findings that perturbations in mechanically active skeletal and cardiac muscle affect function (Zhang et al., 2005, 2010; Lei et al., 2009; Banerjee et al., 2014; Stroud et al., 2017; Zhou et al., 2017; van Ingen & Kirby, 2021). However, while global deletion of multiple LINC components to disrupt the entire complex highlight its importance (Lei et al., 2009; Zhang et al., 2009; Carley et al., 2021), these studies do not reveal functions of individual LINC components in specific tissues. Whether the SUN proteins cell autonomously regulate the vascular endothelium, which is also mechanically active due to outward pressure and shear stress from blood flow, has not been explored.

Mutations in *LMNA* (lamin A/C) cause a premature aging syndrome linked to cardiovascular defects (Capell & Collins, 2006). The LINC complex protein SUN1 is mis-expressed in this disease (Chen et al., 2012), and cellular defects due to the *LMNA* mutation are rescued by reduced levels of SUN1 protein, highlighting a potential function for SUN1 in the disease pathology (Chen et al., 2012; Chang et al., 2019).

Here, we present an in-depth analysis of how the LINC complex component SUN1 affects blood vessel development and function *in vivo*. We found that *Sun1* cell-autonomously regulates blood vessel sprouting and barrier function *in vivo,* and these effects are consistent with SUN1 regulating endothelial cell functions via adherens junction activity. In primary endothelial cells, nuclear SUN1 coordinates peripheral microtubule dynamics that in turn regulate peripheral RhoGEF activation, junction stability and barrier function. Thus, nuclear SUN1 that resides far from endothelial cell junctions regulates endothelial cell-cell communication and blood vessel sprouting via a novel microtubule-based integration pathway from the nucleus to the cell periphery.

## RESULTS

### The nuclear LINC protein SUN1 regulates vascular development

The LINC complex is important for cell migration (Chancellor et al., 2010; King et al., 2014; Denis et al., 2021), and blood vessel formation involves extensive endothelial cell migration; thus, we hypothesized that the LINC complex regulates angiogenic sprouting. Because mutations in endothelial cell *LMNA* causative for human cardiovascular disease are associated with expression changes in the LINC protein SUN1 (Chen et al., 2012), and because *Sun1* has not been functionally analyzed in the vascular endothelium *in vivo*, we first asked whether SUN1 is required for vascular development. Utilizing a mouse line carrying a conditional *Sun1* allele that we generated from *Sun1^tm1a^* “knockout first” mice **(Figure 1-figure supplement 1A-B)**, *Sun1^fl/fl^* mice were bred to *Sun1^fl/+^;Cdh5CreERT2/+* mice to generate *Sun1^iECKO^* (inducible endothelial cell knockout) mice with both endothelial cell-selective and temporal control over *Sun1* excision. Examination of lung DNA, which is rich in endothelial cells, revealed appropriate excision *in vivo* **(Figure 1-figure supplement 1C)**.

The retinal vasculature of *Sun1^iECKO^* pups injected with tamoxifen at P (postnatal day) 1-3 and harvested at P7 had significantly reduced radial expansion relative to littermate controls **(Figure 1A-C, Figure 1-figure supplement 1D)**, consistent with a role for *Sun1* in vascular development. *Sun1^iECKO^* retinas also had increased density at the vascular front **(Figure 1-figure supplement 1E)**, consistent with defects in sprouting that prevent expansion and thus increase density (Hellström et al., 2007; Ricard et al., 2012; Angulo-Urarte et al., 2018). Since vessel densities in the plexus ahead of arteries and veins exhibit heterogeneity, we measured by area and found increased density in the plexus ahead of both arteries and veins in *Sun1^iECKO^* retinas **(Figure 1B, D)**. Because adherens junction dynamics regulate vascular sprouting (Sauteur et al., 2014), this mutant phenotype suggested that endothelial cell-cell junctions were affected by loss of *Sun1*. VE-cadherin localization, a readout of junction integrity (Huveneers et al., 2012; Bentley et al., 2014; Wylie et al., 2018; Vion et al., 2020), was significantly less linear and more punctate in *Sun1^iECKO^* vessels, indicating increased adherens junction turnover and junction instability **(Figure 1E-F)**. Dextran injection was used to functionally evaluate the effects of *Sun1* loss on vascular barrier function *in vivo*, and *Sun1^iECKO^* mice had increased signal in the surrounding tissue compared to controls **(Figure 1-figure supplement 1F)**, suggesting increased vessel permeability. Together, these data indicate a specific role for SUN1 in angiogenic sprouting and endothelial cell-cell junctions *in vivo*.

**Figure 1.**
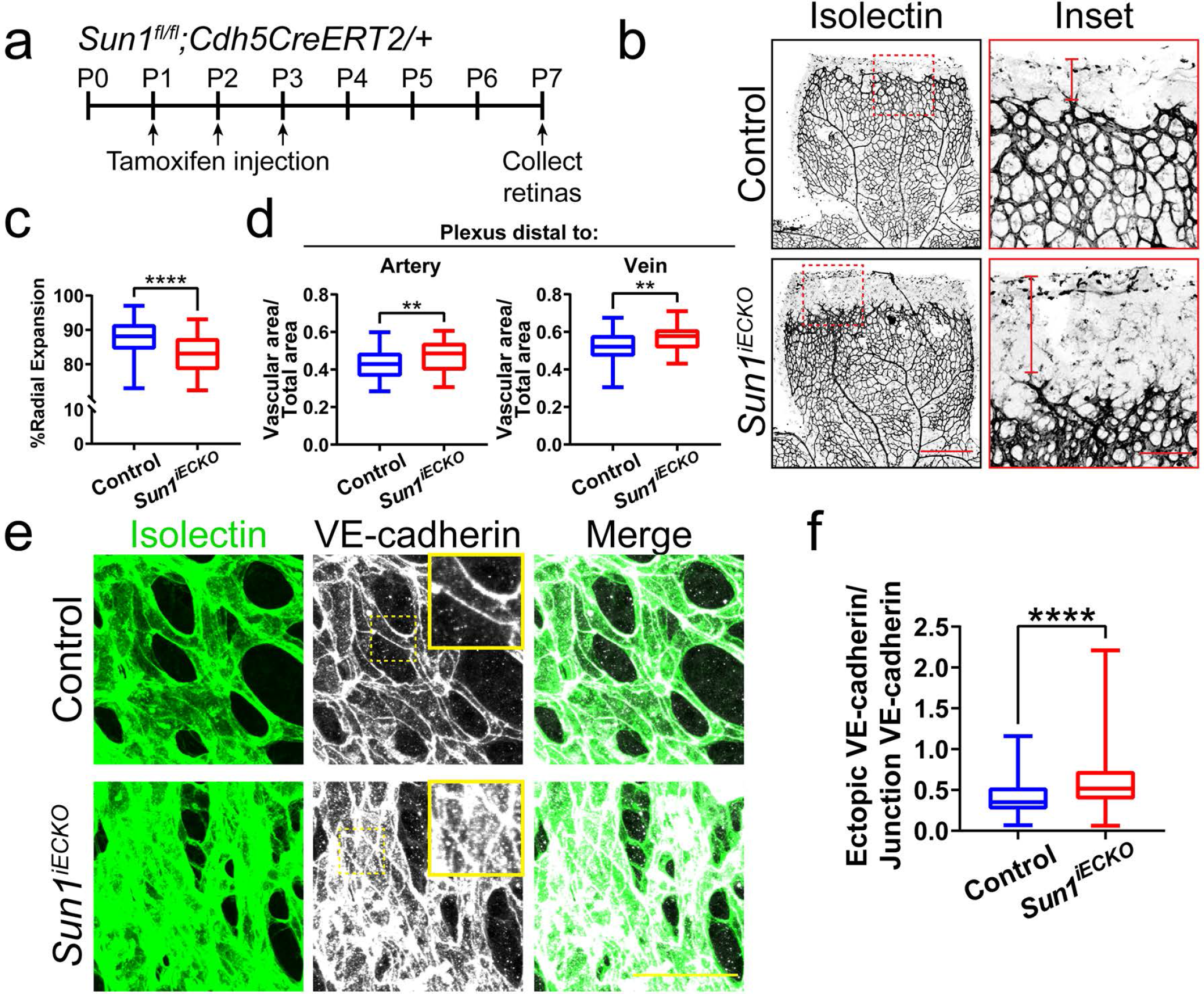
The nuclear LINC protein SUN1 regulates vascular development. **(a)** Schematic of tamoxifen-induced excision of exon 4 of *Sun1* in pups from cross of *Sun1^fl/fl^* X *Sun1^fl/+^;Cdh5CreERT2* mice. **(b)** Representative images of P7 mouse retinas of indicated genotypes, stained for IB4 (isolectin). Scale bar, 500µm. Inset shows vascular plexus ahead of vein. Red line shows expansion of vascular front. Scale bar inset, 150µm. **(c)** Quantification of vessel network radial expansion in **b**. n=186 ROIs from 44 retinas (controls) and 63 ROIs from 16 retinas (*Sun1^iECKO^*) from 6 independent litters. ****, *p*<0.0001 by student’s two-tailed unpaired *t*-test. **(d)** Quantification of vascular density ahead of either arteries or veins. n=87 ROIs (controls, artery), 38 ROIs (*Sun1^iECKO^*, artery), 84 ROIs (controls, vein), and 37 ROIs (*Sun1^iECKO^*, vein) from 27 retinas (controls) and 12 retinas (*Sun1^iECKO^*) from 3 independent litters. **, *p*<0.01 by student’s two-tailed unpaired *t*-test. **(e)** Representative images of IB4 (isolectin) (green, vessels) and VE-cadherin (white, junctions) staining in P7 retinas of indicated genotypes. Scale bar, 50µm. **(f)** Quantification of ectopic VE-cadherin as shown in **e**. n=144 junctions (9 retinas, controls) and 160 junctions (10 retinas, *Sun1^iECKO^*). ****, *p*<0.0001 by student’s two-tailed unpaired *t*-test.

### Nuclear SUN1 is required for sprouting angiogenesis

*Sun1* loss disrupts vascular development in the postnatal mouse retina **(Figure 1)**, but this tissue is not amenable to long-term live image analysis. To query dynamic aspects of angiogenic sprouting, which occurs via regulated changes in endothelial adherens junction stability (Sauteur et al., 2014; Angulo-Urarte et al., 2018; Wylie et al., 2018), we utilized a 3D sprouting model (Nakatsu & Hughes, 2008) coupled with temporal image acquisition. Reduced levels of endothelial cell SUN1 via siRNA knockdown (KD) **(Figure 2-figure supplement 1A-B)** led to significantly decreased sprout length and branching **(Figure 2A-C)**, reminiscent of the decreased radial expansion of *Sun1^iECKO^* retinal vessels described above. SUN1 depletion did not significantly influence the proportion of EdU-labeled or Ki67 stained cells **(Figure 2-figure supplement 1C-F)**, indicating that the abnormal sprouting and branching is not downstream of reduced proliferation. Live-cell imaging revealed that control sprouts typically elongated over time, with little retraction once they extended from the bead **(Figure 2D-E, Movie 1)**. In contrast, SUN1 KD sprouts retracted more often, and many mutant sprouts collapsed partially or completely **(Figure 2D-E, Movie 2)**. SUN1 KD sprouts also showed a more diffuse VE-cadherin junction pattern **(Figure 2F-G)**, similar to those of *Sun1^iECKO^* mice and indicative of over-activated junctions. Thus, SUN1 is required for proper vascular sprout dynamics and morphology, and reduced sprout length and branching are likely downstream of excess sprout retractions and perturbed junctions in SUN1-depleted vessels.

**Figure 2.**
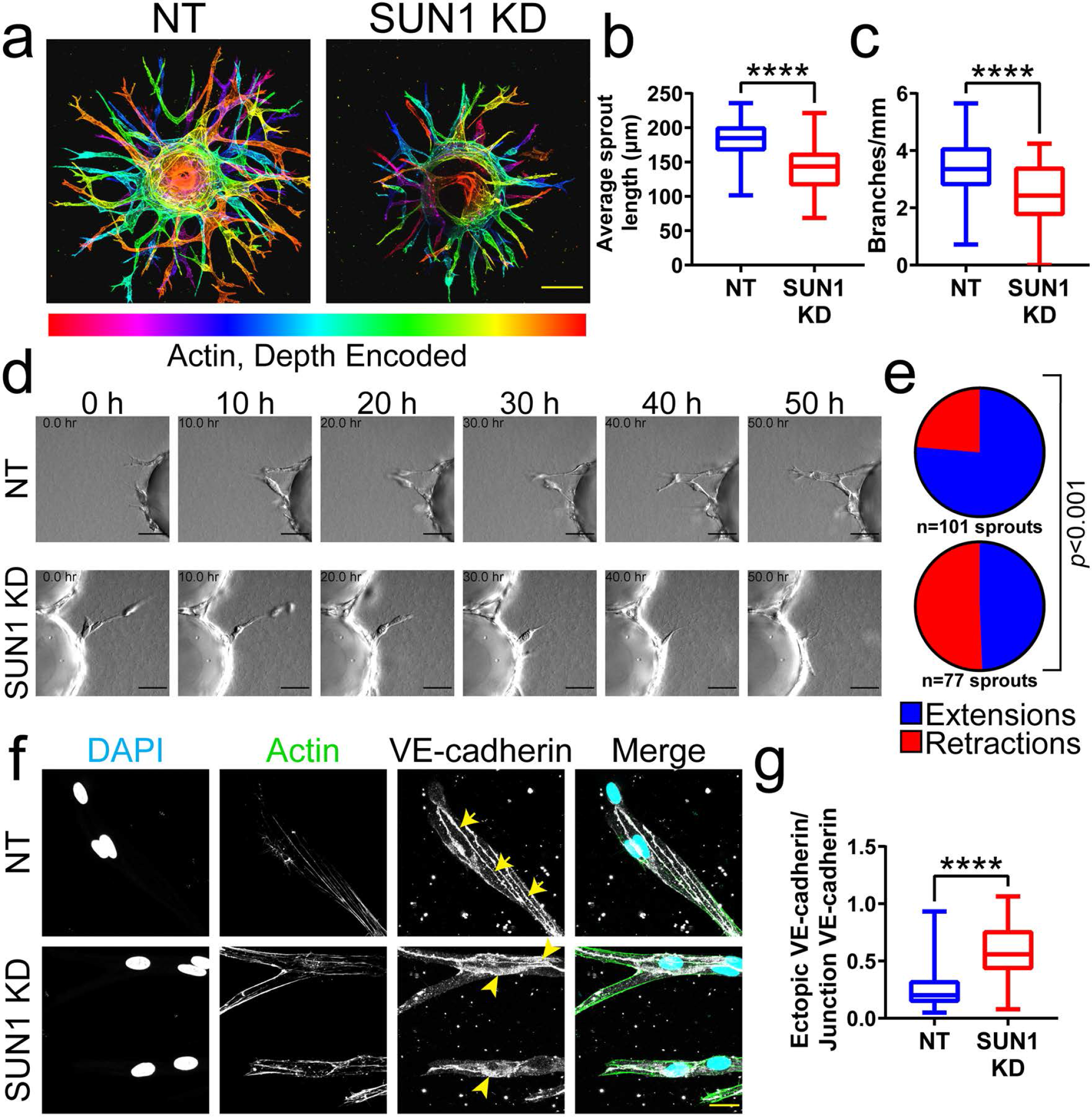
Nuclear SUN1 is required for sprouting angiogenesis. **(a)** Representative images of HUVEC with indicated siRNAs in 3D angiogenic sprouting assay. Sprouts were stained for Phalloidin (actin) and then depth encoded such that cooler colors are further in the Z-plane and warmer colors are closer in the Z-plane. Scale bar, 100µm. **(b)** Quantification of average sprout length of 3D angiogenic sprouts shown in **a**. n=42 beads (NT) and 43 beads (SUN1 KD) compiled from 5 replicates. ****, 3D angiogenic sprouts shown in **a**. n=41 beads (NT) and 43 beads (SUN1 KD) compiled from 5 replicates. ****, *p*<0.0001 by student’s two-tailed unpaired *t*-test. **(d)** Stills from Movie S1 and Movie S2 showing sprouting dynamics of HUVEC with indicated siRNAs over 50h. Scale bar, 50µm. **(e)** Quantification of HUVEC sprout extensions and retractions shown in **d**. n=101 sprouts (NT) and 77 sprouts (SUN1 KD) compiled from 3 replicates. *p*<0.001 by χ^2^ analysis. **(f)** Representative images of HUVEC with indicated siRNAs and stained with indicated antibodies in the 3D sprouting angiogenesis assay. Endothelial cells were stained for DAPI (cyan, DNA), phalloidin (green, actin), and VE-cadherin (white, junctions). Arrows indicate normal junctions; arrowheads indicate abnormal junctions. Scale bar, 20µm. **(g)** Quantification of ectopic VE-cadherin as shown in **f**. n=32 junctions (NT) and 30 junctions (SUN1 KD) compiled from 2 replicates. ****, *p*<0.0001 by student’s two-tailed unpaired *t*-test.

The LINC complex is important for mechanotransduction in muscle fibers with high mechanical loads (van Ingen & Kirby, 2021), and sprouting angiogenesis is regulated by mechanical forces arising from blood pressure and blood flow (Huang et al., 2003). To determine whether SUN1 also regulates sprouting dynamics under laminar flow *in vivo*, we analyzed embryonic zebrafish using a *Tg(fli:LifeAct-GFP)* reporter that labels the endothelial actin cytoskeleton. Zebrafish have two *Sun1* genes, *sun1a* and *sun1b*; the SUN domain of *sun1b* is more homologous to human *SUN1,* and Sun1b is more highly expressed in cardiovascular tissue, so this gene was chosen for manipulation. Sun1b depletion in zebrafish embryos via morpholino (MO) injection led to significantly increased numbers of shorter endothelial cell filopodia at 33-34 hpf (hours post fertilization) in the inter-segmental vessels (ISVs) that sprout towards and connect to the dorsal longitudinal anastomotic vessel (DLAV) **(Figure 3A-C)**. Like the morphants, fish carrying a point mutation in the *sun1b* gene leading to a premature stop codon (*sun1b^sa33109^*, see Methods for details) had shorter filopodia, although filopodia numbers were unchanged in the mutant background **(Figure 3D-F)**. Because increased filopodia are typically seen in actively migrating and sprouting endothelial cells (DeLisser, 2011), these changes are consistent with Sun1b regulating endothelial cell activation in developing zebrafish vessels exposed to physiological flow forces.

**Figure 3.**
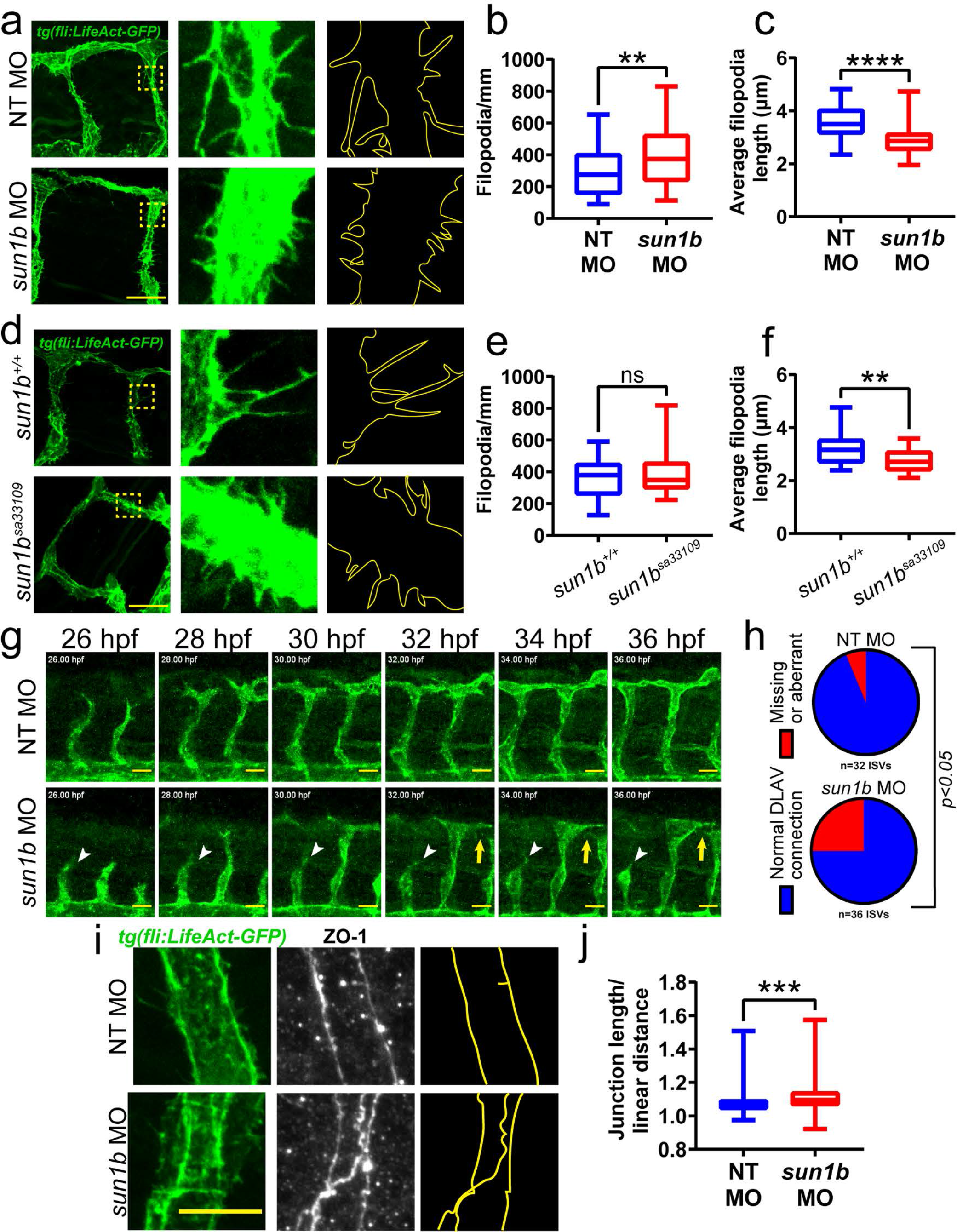
SUN1 regulates actin dynamics and angiogenic sprout extension in 1391 vivo. **(a)** Representative images of zebrafish embryos at 34 hpf with indicated morpholino treatments; anterior to left. *Tg(fli:LifeAct-GFP)* (green, vessels). Insets show ISVs with filopodia, outlines highlighted to show filopodia. Scale bar, 20µm. **(b)** Quantification of filopodia number shown in **a**. n=39 ROIs (15 fish, NT) and 56 ROIs (20 fish, *sun1b* MO) compiled from 3 replicates. **, *p*<0.01 by student’s two-tailed unpaired *t*-test. **(c)** Quantification of average filopodia length shown in **a**. n=39 ROIs (15 fish, NT MO) and 56 ROIs (20 fish, *sun1b* MO) compiled from 3 replicates. ****, *p*<0.0001 by student’s two-tailed unpaired *t*-test. **(d)** Representative images of zebrafish embryos at 34 hpf with indicated genotypes; anterior to left. *Tg(fli:LifeAct-GFP)* (green, vessels). Insets Quantification of filopodia number shown in **d**. n=27 ROIs (9 fish, *sun1b^+/+^*) and 30 ROIs (10 fish, *sun1b ^sa33109^*) compiled from 2 replicates. ns, not significant by student’s two-tailed unpaired *t*-test. **(f)** Quantification of average filopodia length shown in **d**. n=27 ROIs (9 fish, *sun1b ^+/+^*) and 30 ROIs (10 fish, *sun1b ^sa33109^*) compiled from 2 replicates. **, *p*<0.01 by student’s two-tailed unpaired *t*-test. **(g)** Stills from Movie S3 and Movie S4 showing ISV sprouting from 26 hpf to 36 hpf in zebrafish embryos with indicated morpholino treatment; anterior to left. *Tg(fli:LifeAct-GFP)* (green, vessels). White arrowhead points to ISV that does not extend or connect to DLAV. Yellow arrow points to ISV that extends but does not connect to DLAV. Scale bar, 20µm. **(h)** Quantification of ISV connection to DLAV shown in **g**. n=32 ISVs (6 fish, NT MO) and 36 ISVs (6 fish, *sun1b* MO) compiled from 2 replicates. *p*<0.05 by χ^2^ analysis. **(i)** Representative images of zebrafish embryos at 34 hpf with indicated morpholino treatments; anterior to left. *Tg(fli:LifeAct-GFP)* (green, vessels); ZO-1 (white junctions). Outlines highlighted to show junction shapes. Scale bar, 10µm. **(j)** Quantification of junction morphology shown in **i**. n=136 junctions (21 fish, NT MO) and 142 junctions (22 fish, *sun1b* MO). ***, *p*<0.001 by student’s two-tailed unpaired *t*-test.

Next, *sun1b* morphant fish were imaged from 26-36 hpf to determine the effects of Sun1b depletion on vascular sprouting *in vivo*. In controls, the ISVs sprouted towards the dorsal plane and connected to the DLAV between 32-36 hpf **(Figure 3G-H, Movie 3)**. In contrast, numerous ISVs either failed to reach the DLAV or made aberrant connections in *sun1b* morphant fish **(Figure 3G-H, Movie 4)**. Staining for the tight junction protein ZO-1 revealed less linear and more abnormally shaped junctions in *sun1b* morphant fish **(Figure 3I-J)**. These results complement the 3D sprouting analysis and show that the nuclear LINC complex protein SUN1 is important in regulating endothelial cell sprouting dynamics and junction morphology under flow forces *in vivo*.

### SUN1 stabilizes endothelial cell-cell junctions and regulates barrier function

SUN1 loss or depletion in mouse, zebrafish, and 3D sprouting models resulted in abnormal endothelial cell-cell junctions and sprouting behaviors. Thus, we examined more rigorously the hypothesis that SUN1 regulates endothelial cell junction stability and morphology. Primary human umbilical vein endothelial cells (HUVEC) in confluent monolayers had more serrated cell-cell junctions without altered levels of VE-cadherin protein expression after SUN1 depletion, indicative of activated and destabilized junctions **(Figure 4A, Figure 4-figure supplement 1A-B)**. Activated and destabilized endothelial cell adherens junctions are associated with impaired vascular barrier function, so we measured electrical resistance across endothelial monolayers using Real Time Cell Analysis (RTCA) that provides an impedance value positively correlated with barrier function. SUN1 KD endothelial cells had reduced electrical resistance compared to controls **(Figure 4B-C)**, indicative of impaired barrier function and consistent with the increased dextran permeability *in vivo*. Endothelial cells *in vivo* are exposed to blood flow that remodels endothelial junctions (Seebach et al., 2000; Yang et al., 2020). SUN1 depleted HUVEC exposed to laminar shear stress for 72h elongated and aligned properly, but adherens junctions were more serrated and allowed for significantly more matrix exposure as assessed by a biotin labeling assay (Dubrovskyi et al., 2013) under both static and flow conditions **(Figure 4D-F, Figure 4-figure supplement 1C-D)**. These findings support the hypothesis that nuclear SUN1 regulates vascular barrier function via effects on endothelial cell junction stability and morphology.

**Figure 4.**
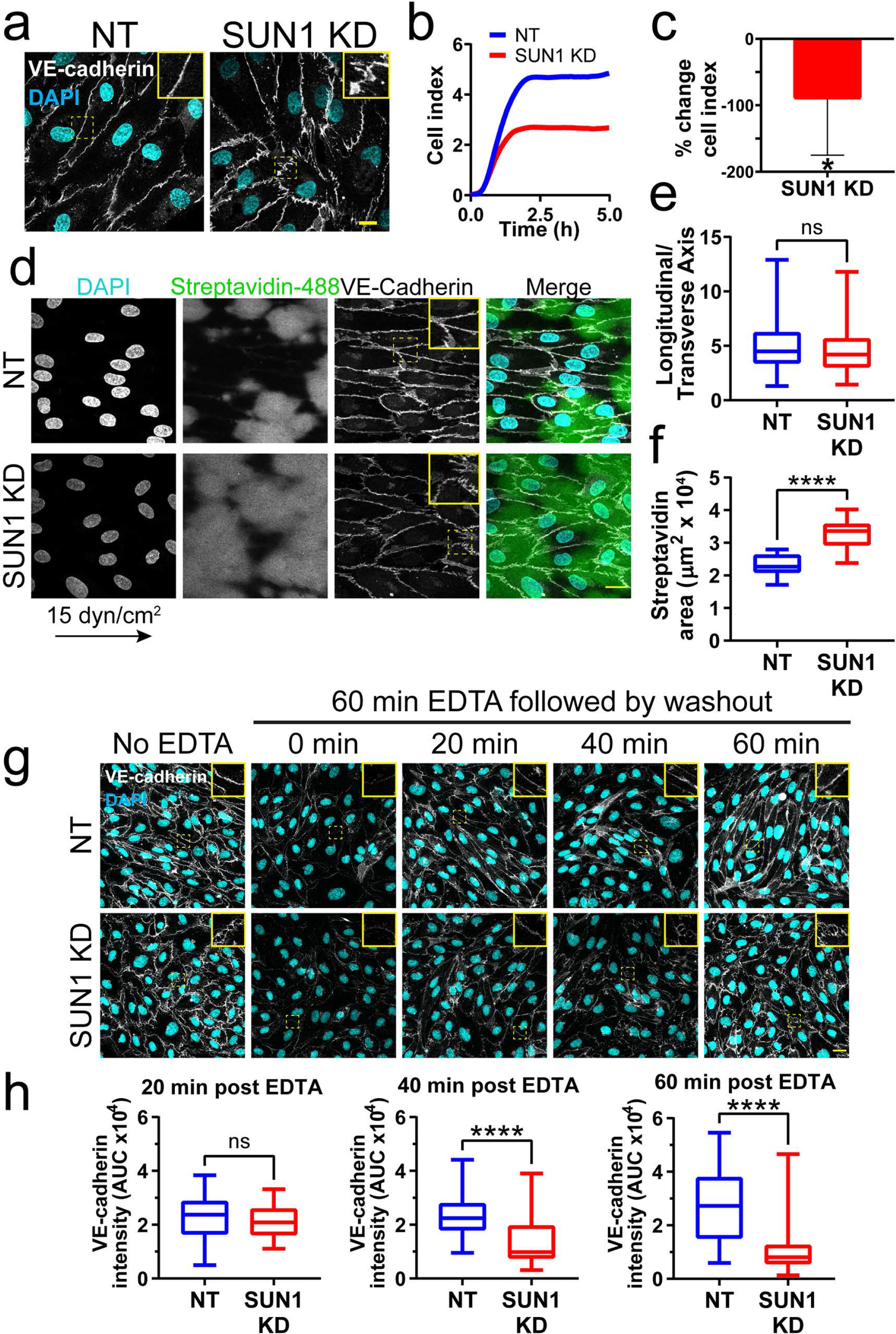
SUN1 stabilizes endothelial cell-cell junctions and regulates barrier 1420 function. **(a)** Representative images of HUVEC with indicated knockdowns in monolayers. Endothelial cells were stained for DAPI (cyan, DNA) and VE-cadherin (white, junctions). Insets show junctions. Scale bar, 10µm. **(b)** Representative graph of impedance measured by RTCA. **(c)** Quantification of % change in cell index for RTCA measured at 5h. Normalized to NT cell index. n=5 replicates. *, *p*<0.05 by student’s two-tailed biotinylated fibronectin and exposed to 15 dyn/cm^2^ shear stress for 72 h then treated with streptavidin. Endothelial cells were stained for DAPI (cyan, DNA), streptavidin (green), and VE-cadherin (white, junctions). Arrow indicates flow direction. Insets show junctions. Scale bar, 20µm. **(e)** Quantification of cell alignment shown in **d**. n=59 cells (NT) and 73 cells (SUN1 KD) compiled from 3 replicates. **(f)** Quantification of streptavidin area shown in **d**. n=15 ROIs (NT) and 15 ROIS (SUN1 KD) compiled from 3 replicates. **(g)** Representative images of HUVEC with indicated siRNAs showing adherens following EDTA washout. Endothelial cells were stained for DAPI (cyan, DNA) and VE-cadherin (white, junctions). Insets show junctions. Scale bar, 20µm. **(h)** Quantification of VE-cadherin line scans at 20 min, 40 min, and 60 min post EDTA washout in **g**. 20 min: n=31 junctions (NT) and 23 junctions (SUN1 KD); 40 min: n=49 junctions (NT) and 33 junctions (SUN1 KD); 60 min: n=33 junctions (NT) and 33 junctions (SUN1 KD) compiled from 3 replicates. ns, not significant; ****, *p*<0.0001 by student’s two-tailed unpaired *t*-test.

Dysfunctional cell-cell junctions can result from abnormal junction formation or the inability of formed junctions to stabilize. To determine how SUN1 functionally regulates endothelial junctions, adherens junctions were disassembled via Ca^2+^ chelation, then reformed upon chelator removal. Junctions were measured using line scans of VE-cadherin intensity along the cell-cell junctions **(Figure 4-figure supplement 1E)**, such that junctions with a linear VE-cadherin signal (stable) had a higher value than those with more serrated patterns (destabilized). No significant difference between SUN1 KD and control junctions was seen at early times post-washout, indicating that SUN1 depletion does not affect adherens junction formation **(Figure 4G-H)**. However, later times post-washout revealed a significant increase in serrated junctions and gaps between endothelial cells in SUN1 KD monolayers relative to controls **(Figure 4G-H)**. Consistent with these findings, SUN1 KD endothelial cells also had increased VE-cadherin internalization at steady state, consistent with actively remodeling junctions **(Figure 4-figure supplement 1F-G)**. Thus, SUN1 is not required to form endothelial cell-cell junctions but is necessary for proper junction maturation and stabilization.

### SUN1 regulates microtubule localization and dynamics in endothelial cells absent effects on gene transcription

We next considered how SUN1 that resides in the nuclear membrane regulates cell behaviors at the cell periphery. SUN1 may affect cell junctions directly via the cytoskeleton, or SUN1 may indirectly affect junctions downstream of gene expression regulation. Because SUN1 is reported to regulate gene transcription and RNA export in other cell types (Li et al., 2017; May & Carroll, 2018), we asked whether endothelial cell transcriptional profiles were altered by SUN1 depletion. To our surprise, RNASeq analysis of HUVEC under both static and flow conditions revealed essentially no significant changes in RNA profiles except for *SUN1* itself **(Table 1, Figure 5-figure supplement 1)**, while in the same experiment we documented extensive expression changes in both up- and down-regulated genes between control HUVEC in static vs. laminar flow conditions, as we and others have shown (Conway et al., 2010; Liu et al., 2021; Maurya et al., 2021; Ruter et al., 2021). These findings are similar to a re-analysis of HeLa cell data (data not shown, (Li et al., 2017)) that revealed few significant changes in gene expression following SUN1 depletion. Thus, although the LINC complex is important for nuclear communication that can affect gene expression, SUN1 is not required for this communication in endothelial cells.

**Table 1.** SUN1 depletion does not alter endothelial gene expression.

| *Condition* | *# Upregulated*<br>*DEG* | *# Downregulated*<br>*DEG* |
| --- | --- | --- |
| *NT_FLOW vs NT_STAT* | 1323 | 1109 |
| *SUN1_STAT vs NT_STAT* | 0 | 1 |
| *SUN1_FLOW vs NT_FLOW* | 1 | 1 |

Abbr: NT, non-targeting; STAT, static; DEG, differentially expressed genes. Red numbers indicate that single downregulated gene was *SUN1*.

We hypothesized that SUN1 directly regulates endothelial cell junction stability via the cytoskeleton. SUN1 has a functional relationship with the microtubule cytoskeleton (Zhu et al., 2017), and microtubule dynamics regulate endothelial cell-cell junctions (Sehrawat et al., 2008, 2011; Komarova et al., 2012). Thus, we asked whether the observed adherens junction defects following SUN1 depletion resulted from changes in the microtubule cytoskeleton. Microtubule depolymerization via nocodazole treatment phenocopied SUN1 KD and destabilized adherens junctions in control monolayers but did not exacerbate the junction defects seen with SUN1 KD **(Figure 5A-B)**. These findings suggest that endothelial cell junction defects are downstream of microtubule perturbations induced by SUN1 depletion.

**Figure 5.**
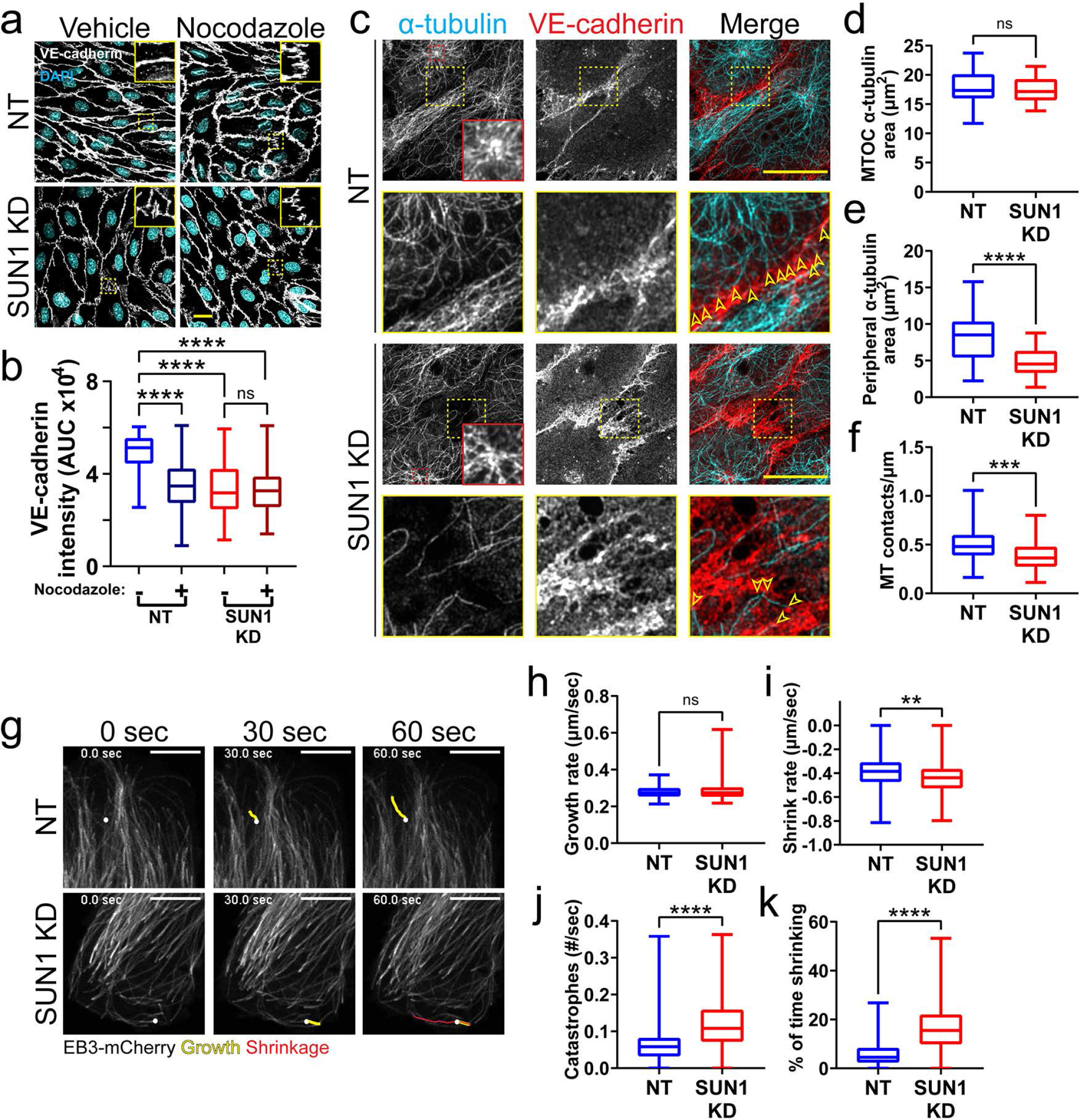
SUN1 regulates microtubule localization and dynamics in endothelial 1443 cells. **(a)** Representative images of HUVEC with indicated siRNAs and indicated treatments. Endothelial cells were stained for DAPI (cyan, DNA) and VE-cadherin (white, junctions). Insets show junctions. Scale bar, 20µm. **(b)** Quantification of VE-cadherin line scans for treatments shown in **a**. n=106 junctions (NT, vehicle), 101 junctions (NT, Nocodazole), 105 junctions (SUN1 KD, vehicle), and 96 junctions (SUN1 KD, Nocodazole) compiled from 3 replicates. ns, not significant; ****, *p*<0.0001 by two-way ANOVA with Tukey’s multiple comparisons test. **(c)** Representative images of HUVEC with indicated siRNAs. junctions). Red insets show α-tubulin at the MTOC (microtubule organizing center), yellow insets show α-tubulin contacts at junctions. Scale bar, 20µm. **(d)** Quantification of α-tubulin area at the MTOC shown in **c**. n=19 cells (NT) and 10 cells (SUN1 KD) compiled from 3 replicates. ns, not significant by student’s two-tailed unpaired *t*-test. **(e)** Quantification of peripheral α-tubulin area shown in **c**. n=39 cells (NT) and 46 cells (SUN1 KD) compiled from 3 replicates. ****, *p*<0.0001 by student’s two-tailed unpaired *t*-test. **(f)** Quantification of contacts between α-tubulin and VE-cadherin shown in **c**. n=75 junctions (NT) and 48 junctions (SUN1 KD) compiled from 3 replicates. ***, *p*<0.001 by student’s two-tailed unpaired *t*-test. **(g)** Stills from Movie S5 and Movie S6 showing microtubule growth in EB3-mCherry labeled HUVEC. White dot indicates start of track. Yellow line indicates growth, red line indicates shrinkage. Scale bar, 10µm. **(h)** Quantification of microtubule growth rate from EB3-mCherry microtubule tracking. N=120 microtubules (12 cells, NT) and 117 microtubules (12 cells, SUN1 KD) compiled from 2 replicates. ns, not significant by student’s two-tailed unpaired *t*-test. **(i)** Quantification of microtubule shrink rate from EB3-mCherry microtubule tracking. n=120 microtubules (12 cells, NT) and 117 microtubules (12 cells, SUN1 KD) compiled from 2 replicates. **, *p*<0.01 by student’s two-tailed unpaired *t*-test. **(j)** Quantification of catastrophe rate from EB3-mCherry microtubule tracking. n=120 microtubules (12 cells, NT) and 117 microtubules (12 cells, SUN1 KD) compiled from 2 replicates. ****, *p*<0.0001 by student’s two-tailed unpaired *t*-test. **(k)** Quantification of percent of time spent shrinking from EB3-mCherry microtubule tracking. n=120 microtubules (12 cells, NT) and 117 microtubules (12 cells, SUN1 KD) compiled from 2 replicates. ****, *p*<0.0001 by student’s two-tailed unpaired *t*-test.

We next investigated how SUN1 affects microtubule localization in endothelial cell monolayers and found that significantly fewer microtubules reached the cell periphery or surrounded the nucleus in SUN1 depleted cells **(Figure 5C, E, Figure 5-figure supplement 2A-B)**. The changes in peripheral microtubule localization were accompanied by significantly fewer microtubule-junction contacts, while α-tubulin levels around the MTOC were not affected **(Figure 5C, D, F)**. Microtubule dynamics were assessed via tip tracking using mCherry-labeled tip protein EB3, which decorated the microtubule lattice and concentrated at growing microtubule tips **(Figure 5G)**. This labeling pattern can occur with EB overexpression but does not affect growth rate (Komarova et al., 2005), so the decorated lattice was used to assess microtubule catastrophe and shrinkage. Microtubules in SUN1 depleted cells had increased shrinkage rates (a more negative value) coupled with increased catastrophe rate and time spent shrinking **(Figure 5G-K, Movies 5, 6)**, consistent with elevated microtubule depolymerization and impaired microtubule dynamics downstream of SUN1 loss, while the microtubule growth rate was unchanged. Taken together, these results indicate that loss of SUN1 impairs microtubule localization and dynamics, and these changes associate with destabilized endothelial cell junctions.

### SUN1 regulates endothelial cell contractility

Microtubule dynamics communicate with the actin cytoskeleton to regulate cell-cell junctions (Verin et al., 2001; Birukova et al., 2004, 2006), and cellular changes in actomyosin contractility contribute to junction activation (Rauzi et al., 2010; Huveneers et al., 2012). We hypothesized that actomyosin mis-regulation induced by SUN1 depletion contributes to endothelial cell junction destabilization and found that SUN1 depletion led to ectopic stress fibers, distinct radial actin bundles at the periphery, and increased phosphorylated myosin light chain (ppMLC), consistent with increased actomyosin contractility **(Figure 6-figure supplement 1A-B)**. Pharmacological blockade of myosin-II ATPase rescued the destabilized cell junctions, ectopic stress fibers and radial actin structures seen with SUN1 depletion **(Figure 6A-B, Figure 6-figure supplement 1C),** suggesting that SUN1 depletion increases contractility. In contrast, thrombin treatment that induces contractility produced over-activated junctions in controls that phenocopied SUN1 depletion but did not further activate junctions of SUN1 KD cells, indicating that SUN1 loss induces a maximal contractile state **(Figure 6C-D)**. Taken together, these data indicate that SUN1 loss results in hypercontractility and destabilized endothelial cell-cell junctions.

**Figure 6.**
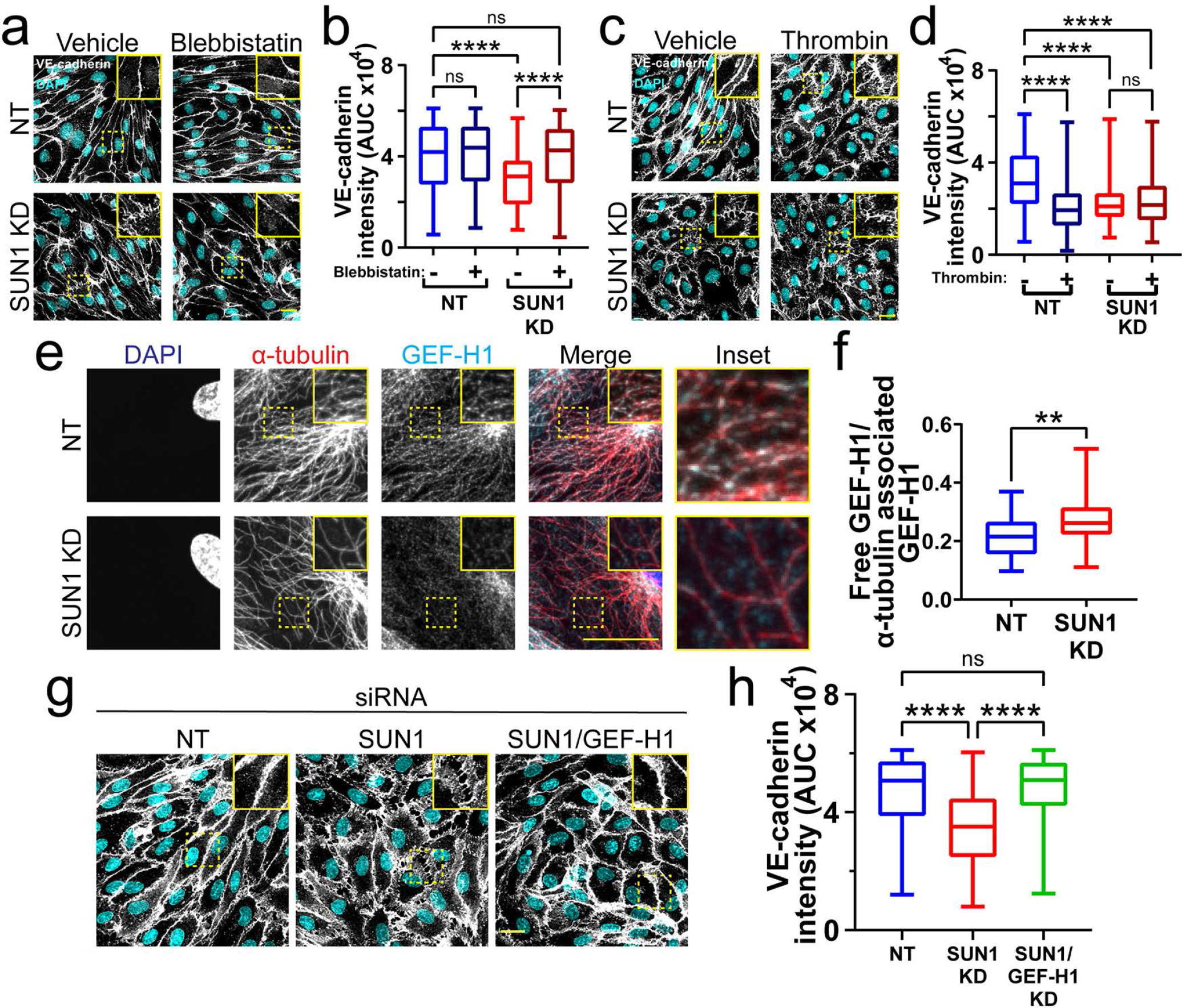
SUN1 regulates endothelial cell contractility and exerts its effects on junctions through the microtubule-associated GEF-H1. **(a)** Representative images of HUVEC with indicated siRNAs and indicated treatments. Endothelial cells were stained for DAPI (cyan, DNA) and VE-cadherin (white, junctions). Insets show junctions. Scale bar, 20µm. **(b)** Quantification of VE-cadherin line scans for treatments shown in **a**. n=159 junctions (NT, vehicle), 154 junctions (NT, blebbistatin), 151 junctions (SUN1 KD, vehicle), and 149 junctions (SUN1 KD, blebbistatin) compiled from 3 replicates. ns, not significant; ****, *p*<0.0001 by two-way ANOVA with Tukey’s multiple comparisons test. **(c)** Representative images of HUVEC with indicated siRNAs and indicated treatments. Endothelial cells were stained for DAPI (cyan, DNA) and VE-cadherin (white, junctions). Insets show junctions. Scale bar, 20µm. **(d)** Quantification of VE-cadherin line scans for treatments shown in **c**. n=75 junctions (NT, vehicle), 70 junctions (NT, thrombin), 71 junctions (SUN1 KD, vehicle), and 73 junctions (SUN1 KD, thrombin) compiled from 3 replicates. ns, not significant; ****, *p*<0.0001 by two-way ANOVA with Tukey’s multiple comparisons test. **(e)** Representative images of HUVEC with indicated siRNAs. Endothelial cells were stained for DAPI (blue, DNA), α-tubulin (red, microtubules), and GEF-H1 (cyan). Insets show α-tubulin and GEF-H1 colocalization. Scale bar, 20µm. **(f)** Quantification of free GEF-H1 normalized to α-tubulin associated GEF-H1 shown in **e**. n=30 cells (NT) and 30 cells (SUN1 KD) compiled from 3 replicates. **, *p*<0.01 by student’s two-tailed unpaired *t*-test. **(g)** Representative images of HUVEC with indicated siRNAs and indicated treatments. Endothelial cells were stained for DAPI (cyan, DNA) and VE-cadherin (white, junctions). Insets show junctions. Scale bar, 20µm. **(h)** Quantification of VE-cadherin line scans from knockdowns shown in **g**. n=169 junctions (NT), 166 junctions (SUN1 KD), 170 junctions (SUN1/GEF-H1 KD) compiled from 3 replicates. ns, not significant; ****, *p*<0.0001 by one-way ANOVA with Tukey’s multiple comparisons test.

### SUN1 affects endothelial junctions through microtubule-associated Rho GEF-H1

To better understand the link between microtubule dynamics, actomyosin contractility, and junction regulation in endothelial cells, we examined Rho signaling downstream of SUN1 in endothelial cells. SUN1 silencing increases RhoA activity in HeLa cells (Thakar et al., 2017), and pharmacological inhibition of Rho kinase (ROCK) signaling in SUN1 depleted endothelial cells rescued the destabilized endothelial cell-cell junctions **(Figure 6-figure supplement 2A-B)** and prevented the radial actin structures **(Figure 6-figure supplement 1D)**. RhoA signaling and barrier function in endothelial cells is regulated by the microtubule-associated RhoGEF, GEF-H1, which is inactive while bound to microtubules and activated following microtubule depolymerization and release (Krendel et al., 2002; Birukova et al., 2006; Birkenfeld et al., 2008). We hypothesized that the impaired microtubule dynamics and Rho-dependent hypercontractility observed following SUN1 depletion were linked via GEF-H1. Consistent with this idea, GEF-H1 strongly localized to peripheral microtubules in control cells but was significantly less localized and more diffuse in SUN1 depleted endothelial cells **(Figure 6E-F)**. Furthermore, depletion of GEF-H1 in endothelial cells that were also depleted for SUN1 rescued the destabilized cell-cell junctions observed with SUN1 KD alone **(Figure 6G-H, Figure 6-figure supplement 2C)**, showing that GEF-H1 is required to transmit the effects of SUN1 depletion to endothelial cell junctions. Thus, nuclear SUN1 normally promotes microtubule-GEFH1 interactions to regulate actomyosin contractility and endothelial cell-cell junction stability, providing a novel linkage from the LINC complex to endothelial cell junctions via the microtubule cytoskeleton.

### SUN1 exerts its effects on endothelial junctions through nesprin-1

Since SUN1 binds nesprins to interact with the cytoskeleton, we considered whether SUN1-nesprin interactions were involved in SUN1 regulation of microtubule dynamics and endothelial cell junction stability. The KASH protein nesprin-1 modulates tight junction protein localization under laminar shear stress (Yang et al., 2020) and regulates microtubule dynamics at the nucleus in muscle syncytia (Gimpel et al., 2017). Co-depletion of SUN1 with nesprin-1 in endothelial cells rescued the effects of SUN1 depletion on junction morphology and functional destabilization measured by matrix biotin labeling **(Figure 7A-B, Figure 7-figure supplement 1A)**. This rescue extended to the cytoskeleton, as co-depletion rescued both decreased peripheral microtubule density and microtubule-GEF-H1 contacts seen with SUN1 depletion **(Figure 7-figure supplement 1B-E)**. Thus, the LINC complex protein nesprin-1 is required to transmit the effects of SUN1 depletion to endothelial cell junctions.

**Figure 7.**
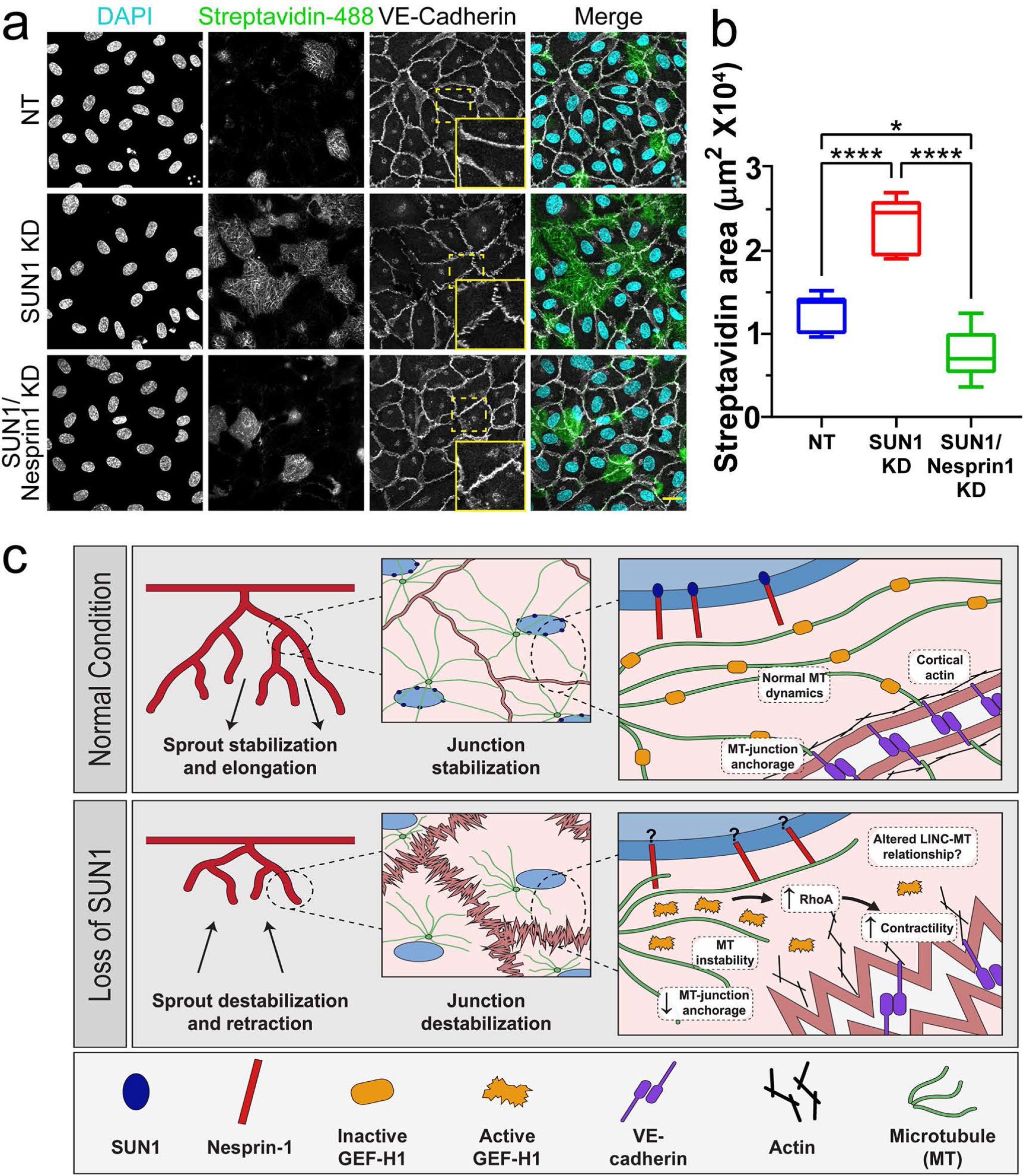
SUN1 regulates endothelial cell junctions through nesprin-1. **(a)** Representative images of HUVEC with indicated siRNAs cultured on biotinylated fibronectin and treated with streptavidin upon confluence. Endothelial cells were stained for DAPI (cyan, DNA), Streptavidin (green), and VE-cadherin (white, junctions). Insets show junctions. Scale bar, 20µm. **(b)** Quantification of streptavidin area shown in **a**. n=6 ROIs (NT), 6 ROIs (SUN1 KD), and 6 ROIs (SUN1/Nesprin-1 KD) from 1 representative replicate. *, *p*<0.05; ****, *p*<0.0001 by one-way ANOVA with Tukey’s multiple comparisons test. **(c)** Model describing proposed role of SUN1 in angiogenic sprouting and endothelial cell junction stabilization.

## DISCUSSION

The nucleus compartmentalizes and organizes genetic material, but how the nucleus directly communicates with other organelles and the cytoskeleton to regulate cell behaviors is poorly understood. Here, we show for the first time that the nuclear LINC complex protein SUN1 regulates angiogenic sprouting and vascular barrier function via long-distance regulation of endothelial cell-cell junctions *in vivo*, and that these effects go through the microtubule cytoskeleton to regulate microtubule dynamics and actomyosin contractility. Our data is consistent with a model in which endothelial SUN1 stabilizes peripheral microtubules that coordinate activity of the microtubule-associated Rho exchange factor GEF-H1. GEF-H1 becomes activated to stimulate Rho signaling upon release from microtubules, and we posit that peripheral microtubules normally regulate local GEF-H1 activity to maintain appropriate microtubule-junction interactions and actomyosin contractility in endothelial cells, leading to cell-cell junctions that remodel to support angiogenic sprouting and maintain vessel barrier integrity **(Figure 7C)**. Loss of SUN1 reduces peripheral microtubules, resulting in GEF-H1 over-activation, elevated RhoA signaling, and increased contractility, leading to destabilized endothelial cell adherens junctions that impair blood vessel formation and function in fish and mice. The LINC complex is further implicated in endothelial junction regulation by our finding that the KASH protein nesprin-1 is required for the effects of SUN1 loss on microtubules, GEF-H1, and proper junction function. Thus, we describe a specific role for SUN1 as a critical mediator of communication between the endothelial cell nucleus and the cell periphery via microtubule regulation of GEF-H1.

We show that the LINC complex protein SUN1 is required for endothelial adherens junction stabilization and proper blood vessel formation and function *in vivo*. Although *Sun2* partially compensates for loss of *Sun1* developmentally (Lei et al., 2009; Zhang et al., 2009) and thus may prevent more profound vascular effects, the significant vascular defects in vessels lacking *Sun1* indicate non-redundant functions for SUN1 in vascular development. The adherens junctions of expanding retinal vessels are destabilized in mice lacking endothelial *Sun1*, and this cellular phenotype is accompanied by increased vessel permeability, reduced radial expansion, and increased network density, indicating that destabilized junctions contribute to the vessel network perturbations. Mutations in other genes that affect endothelial cell junction integrity, such as *Smad6*, *Pi3kca*, and *Yap/Taz*, also perturb retinal angiogenesis (Angulo-Urarte et al., 2018; Neto et al., 2018; Wylie et al., 2018). Live-image analysis of active vessel sprouting showed that SUN1 regulates filopodia dynamics and anastomosis in zebrafish and sprout dynamics in mammalian endothelial cell sprouts, and these vessels also had abnormal junctions in the absence of SUN1. Altered sprout dynamics are found in other scenarios where VE-cadherin is absent or abnormal such as Wnt inhibition, loss of VE-cadherin, PI3-kinase inhibition, and excess centrosomes (Sauteur et al., 2014, 2017; Kushner et al., 2016; Angulo-Urarte et al., 2018; Hübner et al., 2018; Buglak et al., 2020). The LINC complex regulates endothelial cell aggregation into tube-like structures in Matrigel (King et al., 2014; Denis et al., 2021), consistent with our findings that highlight a central role for SUN1 in blood vessel formation *in vivo*. Unlike genes encoding components of junctions and signaling effectors, SUN1 functions at significant cellular distances from cell junctions.

How does SUN1 regulate endothelial cell-cell junctions from a distance? The importance of the LINC complex in transducing signals from the cell periphery and outside the cell to the nucleus to affect nuclear envelope properties and gene transcription is well-studied (Carley et al., 2022); however, how information goes from the nucleus to the cell periphery is less well-understood. Although the LINC complex is clearly important in nucleus-to-cytoplasm communication, most studies have globally manipulated the LINC complex (Zhang et al., 2009; Graham et al., 2018; Carley et al., 2021; Denis et al., 2021), and how individual components contribute is not well-understood. Recently Ueda et al (2022) found that SUN1 regulated focal adhesion maturation in non-endothelial cells *in vitro* via effects on the actin cytoskeleton (Ueda et al., 2022), and we found that SUN1 regulates endothelial cell-cell junctions through microtubules, suggesting that SUN1 is important for signaling from the nucleus to the cell periphery by regulating cytoskeletal organization at several levels. SUN1 is thought to be important in microtubule-associated LINC complex functions (Zhu et al., 2017), and disruption of microtubules or their dynamics destabilizes adherens junctions in both endothelial and non-endothelial cells (Komarova et al., 2012; Vasileva & Citi, 2018).

Microtubule plus end dynamics regulate E-cadherin accumulation at epithelial cell adherens junctions (Stehbens et al., 2006), and dynein is thought to anchor microtubule plus ends to junctions via the plus end binding protein EB1 (Ligon et al., 2001; Shaw et al., 2007; Bellett et al., 2009). Microtubule minus ends also regulate protein accumulation at junctions and are anchored to the junction through a complex involving p120 catenin, PLEKHA7, and CAMSAP3 (Meng et al., 2008). Our data show that SUN1 depletion impairs both microtubule dynamics and microtubule localization. Specifically, elevated rates of microtubule catastrophe and shrinkage absent changes in growth rate likely account for reduced peripheral microtubule density and microtubule-junction contacts. Endothelial junction destabilization was phenocopied by microtubule depolymerization, consistent with our model that the influence of SUN1 on microtubule dynamics and localization is important for the stabilization of endothelial cell junctions.

Here we show that SUN1 regulates endothelial cell junctions via the microtubule-regulated Rho activator GEF-H1. GEF-H1 specifically modulates RhoA signaling at endothelial cell adherens junctions to influence VE-cadherin internalization (Juettner et al., 2019). Microtubule depolymerization releases GEF-H1 to activate RhoA signaling and elevate actomyosin contractility (Krendel et al., 2002; Birkenfeld et al., 2008) in non-endothelial cells, while direct manipulation of GEF-H1 via depletion or blockade attenuates agonist-induced endothelial barrier dysfunction (Birukova et al., 2006). Our work revealed that peripheral GEF-H1 localization was more diffuse following endothelial SUN1 silencing, suggesting its activation with SUN1 loss, and Rho-kinase inhibition rescued the SUN1 depletion-induced destabilization of endothelial cell junctions. These findings support that altered microtubule dynamics downstream of SUN1 depletion promote the release and over-activation of peripheral GEF-H1, and this activation destabilizes cell junctions.

Thus, SUN1 regulation of microtubule dynamics is linked to its regulation of endothelial cell junction stability, although exactly how SUN1 influences microtubule dynamics and function is unclear. SUN1 resides in the nuclear envelope and alters gene transcription in other cell types (Li et al., 2017; May & Carroll, 2018). Our transcriptional analysis of endothelial cells under flow did not reveal significant transcriptome changes with SUN1 depletion, indicating that SUN1 regulation of endothelial junctions occurs via its role in LINC complex interactions with the cytoskeleton. Nesprin-1, a KASH protein that functions in LINC complex-cytoskeletal interactions, is shown here to mediate the effects of SUN1 loss on endothelial junctions, suggesting that SUN1 normally sequesters nesprin-1 to prevent formation of ectopic complexes. Since fewer microtubules surrounded the nucleus in SUN1-depleted endothelial cells, SUN1 may normally prevent abnormal or unstable nesprin-1 LINC complexes that promote microtubule depolymerization, and SUN1 loss allows these complexes to form. SUN1 may compete for nesprin-1 binding with other nuclear envelope proteins that interact with nesprin-1, such as SUN2 or nesprin-3 (Stewart-Hutchinson et al., 2008; Taranum et al., 2012; Yang et al., 2020). This idea is consistent with the finding that SUN1 antagonizes SUN2-based LINC complexes that promote RhoA activity in HeLa cells, although microtubule localization changes were not reported (Thakar et al., 2017).

Our finding that SUN1 regulates endothelial cell barrier function and blood vessel sprouting has implications for diseases associated with aging, as vascular defects underlie most cardiovascular disease. Children with Hutchinson-Gilford Progeria Syndrome (HGPS) have a mutation in the *LMNA* gene encoding lamin A/C, resulting in accumulation of an abnormal lamin protein called progerin; these patients age rapidly and die in their early to mid-teens from severe atherosclerosis (De Sandre-Giovannoli, 2003; Eriksson et al., 2003; Olive et al., 2010). Progerin has increased SUN1 affinity that leads to SUN1 accumulation in HGPS patient cells (Haque et al., 2010; Chen et al., 2012, 2014; Chang et al., 2019). Endothelial cells also accumulate SUN1 in HGPS mouse models (Osmanagic-Myers et al., 2019), and loss of *Sun1* partially rescues progeria phenotypes in mouse models and patient cells (Chen et al., 2012; Chang et al., 2019). Thus, nuclear membrane perturbations affecting SUN1 cause disease, and here we find that nuclear SUN1 regulates microtubules to affect both the microtubule and actin cytoskeletons. These effects are transmitted to endothelial cell-cell junctions far from the site of SUN1 localization to influence endothelial cell behaviors, blood vessel sprouting and barrier function.

## MATERIALS & METHODS

### Microscopy

Unless otherwise stated, all imaging was performed as follows: confocal images were acquired with an Olympus confocal laser scanning microscope and camera (Fluoview FV3000, IX83) using 405nm, 488nm, 561nm, and 640nm lasers and a UPlanSApo 40x silicone-immersion objective (NA 1.25), UPlanSApo 60x oil-immersion objective (NA 1.40), or UPlanSApo 100x oil-immersion objective (NA 1.40). Imaging was performed at RT for fixed samples. Images were acquired with the Olympus Fluoview FV31S-SW software and all image analysis, including Z-stack compression, was performed in Fiji (Linkert et al., 2010; Schindelin et al., 2012). Any adjustments to brightness and contrast were performed evenly for images in an experiment.

### Mice

All experiments involving animals were performed with approval from the University of North Carolina, Chapel Hill Institutional Animal Care and Use Committee (IACUC). C57Bl6N-*Sun1^tm1a(EUCOMM)Wtsi^*/CipheOrl mice were obtained from the European Mouse Mutant Archive (EMMA) mouse repository. FlpO-B6N-Albino (Rosa26-FlpO/+) mice were obtained from the UNC Animal Models Core. *Tg(Cdh5-cre/ERT2)1Rha* mice were generated by Dr. Ralf Adams (Sörensen et al., 2009) and obtained from Cancer Research UK. The *Sun1^tm1a^* allele was identified via genomic PCR to amplify the LacZ insertion (Forward: 5’-ACTATCCCGACCGCCTTACT-3’; Reverse: 5’-TAGCGGCTGATGTTGAACTG-3’). The *Sun1^fl^* allele was generated by breeding *Sun1^tm1a^* mice with FlpO-B6N-Albino (Rosa26-FlpO/+) mice to excise the *lacZ* insertion. The *Sun1^fl^* allele was identified via genomic PCR using the following primers (Forward: 5’-GCTCTCTGAAACATGGCTGA-3’; Reverse: 5’-ATCCGGGGTGTTTGGATTAT-3’). *Sun1^fl^* mice were bred to *Tg(Cdh5-cre/ERT2)1Rha* mice to generate *Sun1^fl/fl^*;*Cdh5CreERT2* pups for endothelial-selective and temporally controlled deletion of exon 4 of the *Sun1* gene. The excised *Sun1* allele was identified via genomic PCR on lung tissue using the following primers (Forward: 5’-CTTTTGGGCTGCTCTGTTGT-3’; Reverse: 5’-ATCCGGGGTGTTTGGATTAT-3’). PCR genotyping for FlpO and Cdh5CreERT2 mice was performed with the following primers (FlpO: Forward: 5’-TGAGCTTCGACATCGTGAAC -3’; Reverse: 5’-TCAGCATCTTCTTGCTGTGG-3’) (Cdh5CreERT2: Forward: 5’-TCCTGATGGTGCCTATCCTC-3’; Reverse: 5’-CCTGTTTTGCACGTTCACCG-3’). Induction of Cre was performed via IP injection of pups at P1, P2, and P3 with 50µl of 1mg/ml tamoxifen (T5648, Sigma) dissolved in sunflower seed oil (S5007, Sigma). Littermates lacking either *Cdh5CreERT2* or the *Sun1^fl^* allele were used as controls.

### Mouse retinas

Tamoxifen-injected mice were sacrificed at P7, eyes were collected, fixed in 4% PFA for 1h at RT, then dissected and stored at 4°C in PBS for up to 2 weeks (Chong et al., 2017). Retinas were permeabilized in 0.5% Triton X-100 (T8787, Sigma) for 1h at RT, blocked for 1h at RT in blocking solution (0.3% Triton X-100, 0.2% BSA (A4503, Sigma), and 5% goat serum (005-000-121, Jackson Immuno)), then incubated with VE-cadherin antibody (anti-mouseCD144, 1:100, 550548, BD Pharmingen) in blocking solution overnight at 4°C. Samples were washed 3X, then incubated with Isolectin B4 AlexaFluor 488 (1:100, I21411, ThermoFisher) and goat anti-rat AlexaFluor 647 (1:500, A21247, Life Technologies) for 1h at RT. Retinas were mounted with Prolong Diamond Antifade mounting medium (P36961, Life Technology) and sealed with nail polish. Images were obtained using either a UPlanSApo 10x air objective (NA 0.40) or UPlanSApo 40x silicone-immersion objective (NA 1.25). Percent radial expansion was calculated by dividing the distance from the retina center to the vascular front by the distance from the retina center to the edge of the tissue. Four measurements/retina were averaged. Vascular density was measured by imaging a 350µm x 350µm ROI at the vascular edge. Fiji was used to threshold images, and the vessel area was normalized to the area of the ROI (n=4 ROI/retina, chosen at 2 arteries and 2 veins). Junctions were measured by taking the ratio of the mean intensity of the junction and the mean intensity of the area immediately adjacent to the junction. Mean intensity was measured via line scans in Fiji. 16 junctions/retina were measured.

### Retina blood vessel permeability

Tamoxifen-injected mice were anesthetized at P7 in isoflurane for 5 min. The abdomen was opened, and the diaphragm was cut. 100µl of 5mg/ml 10kDa Dextran-Texas Red (D1863, Invitrogen) in PBS was injected into the left ventricle of the heart. Eyes were immediately collected and fixed in 4% PFA for 1h at RT, then dissected and stained as described above. Leak was determined by making a mask of the vessel area using the isolectin channel, then assessing the dextran signal outside the vessel.

### 3D sprouting angiogenesis assay

The 3D sprouting angiogenesis assay was performed as previously described (Nakatsu & Hughes, 2008; Nesmith et al., 2017). 48h following siRNA knockdown, HUVEC were coated onto cytodex 3 microcarrier beads (17048501, GE Healthcare Life Sciences) and embedded in a fibrin matrix by combining 7µl of 50U/ml thrombin (T7201-500UN, Sigma) with 500µl of 2.2 mg/ml fibrinogen (820224, Fisher) in a 24-well glass-bottomed plate (662892, Grenier Bio). The matrix was incubated for 20 min at RT followed by 20 min at 37°C to allow the matrix to solidify. EGM-2 was then added to each well along with 200µl of normal human lung fibroblasts (CC2512, Lonza) at a concentration of 2×10^5^ cells/ml. At day 7 of sprouting, fibroblasts were removed via trypsin treatment (5X-trypsin for 3min at 37°C), and samples were fixed in 4% PFA for 15 min at RT. 0.5% Triton X-100 in DBPS was added to the wells, and incubation was overnight at 4°C. After rinsing 3X in DPBS, samples were blocked (5% goat serum (005-000-121, Jackson Immuno), 1% BSA (A4503, Sigma), and 0.3% Triton X-100 (T8787, Sigma)) overnight at 4°C. Samples were rinsed 3X in DPBS then anti-VE-cadherin antibody (1:1000, 2500S, Cell Signaling) in blocking solution was added for 24h at 4°C. Samples were rinsed 3X 10 min in 0.5% Tween 20 then washed overnight at 4°C in 0.5% Tween. Samples were rinsed 3X in DPBS, then DAPI (0.3µM, 10236276001, Sigma) and AlexaFluor488 Phalloidin (1:50, A12379, Life Technologies) in blocking solution were added to the wells, and incubation was overnight at 4°C prior to rinsing 3X in DPBS. For whole bead analysis, images were acquired in the Z-plane using a UPlanSApo 20x oil-immersion objective (NA 0.58) and processed in Fiji.

Average sprout length was measured by tracing each sprout from base (bead) to tip, then averaging lengths per bead. Branching was measured by counting total branch points and normalizing to total sprout length per bead using the AnalyzeSkeleton plugin (Arganda-Carreras et al., 2010). For junctions, images were acquired in the Z-plane using a UPlanSApo 60x oil-immersion objective (NA 1.40). Junctions were measured by taking the ratio of the mean intensity of the junction and the mean intensity of the area immediately adjacent to the junction. Mean intensity was measured via line scans in Fiji.

Live imaging on HUVEC sprouts was performed between days 4-6.5 of sprouting. Images were acquired on an Olympus VivaView Incubator Fluorescence Microscope with a UPLSAPO 20x objective (NA 0.75) and 0.5x magnification changer (final magnification 20x) with a Hamamatsu Orca R2 cooled CCD camera at 30 min intervals for 60h at 37°C. Images were acquired using the MetaMorph imaging software and analyzed in Fiji. Sprouts were considered to have “retracted” if they regressed towards the bead for at least 3 imaging frames (1.5h).

### Zebrafish

All experimental Zebrafish (*Danio rerio*) procedures performed in this study were reviewed and approved by the University of North Carolina Chapel Hill Animal Care and Use Committee. Animals were housed in an AAALAC-accredited facility in compliance with the *Guide for the Care and Use of Laboratory Animals* as detailed on protocols.io (dx.doi.org/10.17504/protocols.io.bg3jjykn). *Tg(fli:LifeAct-GFP)* was a gift from Wiebke Herzog. *sun1b^sa33109^* mutant fish were obtained from the Zebrafish International Resource Center (ZIRC). For genotyping, the target region of the *sun1b* gene was amplified via genomic PCR using the following primers (Forward: 5’-GGCTGCGTCAGACTCCATTA-3’; Reverse: 5’-TTGAGTTAAACCCAGCGCCT-3’). The amplicon was then sequenced by Sanger sequencing (GENEWIZ) using the forward primer. Morphant fish were obtained by injecting 2.5-5ng of non-targeting (NT) (5’-CCTCTTACCTCAGTTACAATTTATA-3’, GeneTools, LLC) or *sun1b* (5’-CGCAGTTTGACCATCAGTTTCTACA-3’, GeneTools, LLC) morpholinos into *Tg(fli:LifeAct-GFP)* embryos at the 1-cell stage. Fish were grown in E3 medium at 28.5°C to 33-34 hpf.

### Zebrafish imaging

Dechorionated embryos were incubated in ice cold 4% PFA at 4°C overnight or RT for 2h. Embryos were permeabilized in 0.5% Triton X-100 in PBST (DPBS + 0.1% Tween 20) for 1 h at RT then blocked (PBST + 0.5% Triton X-100 + 1% BSA + 5% goat serum + 0.01% sodium azide) for 2h at RT. Anti-ZO1 primary antibody (1:500, 33-9100, Thermo Fisher) was added overnight at 4°C. Embryos were rinsed in PBST overnight at 4°C. Goat anti-mouse AlexaFluor647 secondary antibody (1:1000, A-21236, Life Technologies) was added overnight at 4°C. Embryos were washed 3X in PBST for 30 min then overnight in PBST at 4°C. Embryos were rinsed in PBS and a fine probe was used to de-yolk and a small blade to separate the trunk from the cephalic region.

Samples were mounted using Prolong Diamond Antifade mounting medium (P36961, Life Technology) and the coverslip was sealed with petroleum jelly. Imaging was at RT using a UPlanSApo 20x oil-immersion objective (NA 0.58) or a UPlanSApo 60x oil-immersion objective (NA 1.40) with an additional 3x magnification, for a total magnification of 180x. Filopodia length was measured by drawing a line from the filopodia base to the tip. Filopodia number was measured by counting the number of filopodia and normalizing to the total vessel length. Filopodia in at least 3 areas per fish were measured. Junctions were analyzed by drawing a line along the junction then normalizing to the shortest distance between the two ends of the junction. At least 3 junctions were measured per fish.

For live imaging of zebrafish, fish were dechorionated and then anesthetized with 1x Tricaine in E3 for 5 min. Fish were embedded in 0.5% agarose in 1x Tricaine-E3 medium on a glass-bottomed plate in a stage-top incubator (TOKAI HIT, WSKM) at 28.5°C. Images were acquired using a UPlanSApo 40x air objective (NA 0.95) every 15min for 10-15h. ISVs that did not reach the DLAV or connected at non-consistent intervals were considered to have a missing or aberrant DLAV connection if at least 1 ISV posterior to the scored ISV made a normal connection.

### Cell culture

HUVEC (C2519A, Lonza) were cultured in EBM-2 (CC-3162, Lonza) supplemented with the Endothelial Growth Medium (EGM-2) bullet kit (CC-3162, Lonza) and 1x antibiotic-antimycotic (Gibco). Normal human lung fibroblasts (CC2512, Lonza) were cultured in DMEM (Gibco) with 10% fetal bovine serum (FBS) and 1x antibiotic-antimycotic (Gibco). All cells were maintained at 37°C and 5% CO_2_. For contractility inhibition experiments, HUVEC were treated with 10µM (-) Blebbistatin (B0560-1MG, Sigma) for 15 min at 37°C or 10µM Y-27632 (10187-694, VWR) for 30 min at 37°C then immediately fixed in 4% PFA. For induction of contractility, HUVEC were treated with 0.5U/ml thrombin (T7201-500UN, Sigma) for 10 min at 37°C. For microtubule depolymerization, HUVEC were treated with 10µM nocodazole (M1404, Sigma) for 20 min at 37°C then immediately fixed in methanol.

### siRNA knockdown

HUVEC were transfected with non-targeting siRNA (NT, #4390847, Life Technologies), SUN1 siRNA #1 (439240, #s23630, Life Technologies), SUN1 siRNA #2 (439240, #s23629, Life Technologies), GEF-H1 siRNA (439240, #s17546, Life Technologies), and/or nesprin-1 siRNA (M-014039-02-0005, Dharmacon) using Lipofectamine 2000 (11668027, Invitrogen) or Lipofectamine 3000 (L3000015, ThermoFisher). siRNA at 0.48µM in Opti-MEM (31985-070, Gibco) and a 1:20 dilution of Lipofectamine in Opti-MEM were incubated separately at RT for 5 min, then combined and incubated at RT for 15 min. HUVEC were transfected at ∼80% confluency with siRNA at 37°C for 24h, then recovered in EGM-2 for an additional 24h. HUVEC were seeded onto glass chamber slides coated with 5µg/ml fibronectin (F2006-2MG, Sigma) for experiments.

### RNA sequencing and analysis

RNA was extracted using TRIzol (15596018, Invitrogen) from 3 biological replicates (independent experiments) of HUVEC under static or homeostatic laminar flow (15d/cm^2^, 72hr) conditions, and KAPA mRNA HyperPrep Kit (7961901001, Roche) was used to prepare stranded libraries for sequencing (NovaSeq S1). 2–3 × 10^7^ 50-bp paired-end reads per sample were obtained and mapped to human genome GRCh38 downloaded from https://support.10xgenomics.com/single-cell-gene-expression/software/pipelines/latest/advanced/references with STAR using default settings (Dobin et al., 2013). Mapping rate was over 80% for all samples, and gene expression was determined with Htseq-count using the union mode (https://htseq.readthedocs.io/en/master/) (Putri et al., 2022). Differential expression analysis was performed with DESeq2 (Love et al., 2014) using default settings in R, and lists of differentially expressed genes were obtained (p adjusted < 0.1).

### Immunofluorescence staining

For experiments visualizing microtubules, HUVEC were fixed in ice-cold methanol for 10 min at 4°C. For all other experiments, HUVEC were fixed with 4% PFA for 10 min at RT and permeabilized with 0.1% Triton X-100 (T8787, Sigma) for 10 min at RT. Fixed HUVEC were blocked for 1h at RT in blocking solution (5% FBS, 2X antibiotic-antimycotic (Gibco), 0.1% sodium azide (s2002-100G, Sigma) in DPBS). Cells antibiotic-antimycotic (Gibco), 0.1% sodium azide (s2002-100G, Sigma) in DPBS). Cells were incubated in primary antibody overnight at 4°C, then washed 3X for 5 min in DPBS. Secondary antibody and DR (1:1000, ab109202, Abcam), DAPI (0.3µM, 10236276001, Sigma), and/or AlexaFluor488 Phalloidin (1:100, A12379, Life Technologies) were added for 1h at RT followed by 3X washes for 10 min each in DPBS. Slides were mounted with coverslips using Prolong Diamond Antifade mounting medium (P36961, Life Technology) and sealed with nail polish. Primary and secondary antibodies were diluted in blocking solution. The following primary antibodies were used: anti-VE-cadherin (1:500, 2500S, Cell Signaling), anti-SUN1 (1:500, ab124770, Abcam), anti-Ki67 (1:500, ab15580, Abcam), anti-phospho-myosin light chain 2 (Thr18/Ser19) (1:500, 3674S, Cell Signaling), anti-alpha-tubulin (1:500, 3873S, Cell Signaling), anti-GEF-H1 (1:500, ab155785, Abcam), and anti-SYNE1 (1:500, HPA019113, Atlas Antibodies). The following secondary antibodies from Life Technologies were used: goat anti-mouse AlexaFluor 488 (1:500, A11029), goat anti-rabbit AlexaFluor 594 (1:500, A11037), goat anti-mouse 647 (1:500, A21236), and goat anti-rabbit 647 (1:500, A21245).

### Western blotting

Cells were scraped into RIPA buffer with protease/phosphatase inhibitor (5872S, Cell Signaling) then centrifuged at 13000 rpm at 4°C for 20 min. Lysate was reduced in sample loading buffer and dithiothreitol (R0861, Thermo Fisher) and boiled for 10 min at 100°C. Samples were stored at -20°C until use. 10µg of sample were run on a 10% stain-free polyacrylamide gel (161-0183, BioRad) then transferred onto a PVDF membrane on ice for 1.5h. Membranes were blocked in OneBlock (20-313, Prometheus) for 1 h at RT then washed 3X in PBST. Anti-GEF-H1 (1:1000, ab155785, Abcam), anti-VE-cadherin (1:14,000, 2500S, Cell Signaling), or anti-GAPDH (1:5000, 97166S, Cell Signaling) was added overnight at 4°C. Membranes were washed 3X in PBST then donkey anti-rabbit HRP secondary antibody (1:10,000, A16035, Thermo Fisher) was added for 1h at RT. Immobilon Forte HRP Substrate (WBLUF0100, Millipore Sigma) was added for 30 sec. Blots were exposed for 8 sec.

### EdU labeling

HUVEC were labeled with EdU using the Click-It EdU Kit 488 (Invitrogen, C10337) and fixed according to the manufacturer’s instructions. Cells positive for EdU labeling were counted and compared to total cell number to obtain percent positive.

### Junction analysis

Endothelial cell adherens junctions were quantified in monolayers using Fiji to generate 15µm line scans of VE-cadherin signal parallel to the cell junctions. VE-cadherin signal was integrated to obtain the area under the curve. Linear junctions with consistent VE-cadherin signal thus had a large area under the curve, while more serrated junctions had reduced area under the curve **(Fig S3e)**. Measurements were performed on at least 9 cells per field of view, with 3-6 fields of view per condition.

### Real time cell analysis (RTCA)

Barrier function was assessed using the xCELLigence Real-Time Cell Analyzer (RTCA, Acea Biosciences/Roche Applied Science) to measure electrical impedance across HUVEC monolayers seeded onto microelectrodes. HUVEC were seeded to confluency on the microelectrodes of the E-plate (E-plate 16, Roche Applied Science). Electrical impedance readings were taken every 2 min for 5h. The percent change in cell index was obtained at the 5h timepoint using the following formula: (Cell Index_SUN1_- Cell Index_NT_)/ABS(Cell Index_NT_).

### Flow experiments

Flow experiments were performed using an Ibidi pump system (10902, Ibidi) as described (Ruter et al., 2021). HUVEC were seeded onto fibronectin coated Ibidi slides (either µ-Slide I Luer I 0.4 mm (80176, Ibidi) or µ-Slide Y-shaped (80126, Ibidi) in flow medium (EBM-2 with 10% FBS and 1x Antibiotic-antimycotic). HUVEC were exposed to laminar shear stress for 30 min at 5 dyn/cm2, followed by 30 min at 10 dyn/cm2, and finally for 72 h at 15 dyn/cm2. Alignment was measured by taking the ratio of the longitudinal axis to the transverse axis relative to the flow vector. At least 10 cells were measured per condition per experiment. Vascular permeability *in vitro* was determined using the biotin matrix labeling assay as described below.

### Biotin matrix labeling assay

Labeling of biotinylated matrix was assessed as described (Dubrovskyi et al., 2013). Briefly, fibronectin was biotinylated by incubating 0.1mg/mL fibronectin with 0.5mM EZ-Link Sulfo-NHS-LC-Biotin (A39257, ThermoFisher) for 30 min at RT. Glass chamber slides were coated with 5µg/ml biotinylated-fibronectin and HUVEC were seeded on top. At confluency, HUVEC were treated with 25µg/mL Streptavidin-488 (S11223, Invitrogen) for 3 min then immediately fixed. For quantification, Fiji was used to threshold the streptavidin signal, and the streptavidin area was measured and normalized for total area for at least 3 fields of view per experiment.

### Junction reformation assay

The EDTA junction reformation assay was performed as previously described (Wright et al., 2015). Briefly, HUVEC were treated with 3mM EDTA (EDS-100G, Sigma-Aldrich) for 1h at 37°C. EDTA was then washed out 3X with DPBS, incubation at 37°C in EGM-2 was continued, and cells were fixed at 0 min, 20 min, 40 min, and 60 min intervals.

### VE-cadherin internalization

VE-cadherin internalization was performed as described (Wylie et al., 2018). Briefly, HUVEC were plated on 5µg/ml fibronectin and grown to confluency. After overnight serum starvation (Opti-MEM (31985-070, Gibco) supplemented with 1% FBS (F2442, Sigma), and 1x antibiotic-antimycotic (Gibco)), cells were washed with pre-chilled PBS+ (14040182, ThermoFisher) on ice at 4°C, then incubated in ice-cold blocking solution (EBM-2 (CC-3162, Lonza) supplemented with 0.5% BSA (A4503, Sigma)) for 30 min at 4°C. HUVEC were then incubated with VE-cadherin BV6 antibody (1:100, ALX-803-305-C100, Enzo) in blocking solution for 2h on ice at 4°C. Following VE-cadherin labeling, cells were washed with PBS+ then incubated in pre-warmed internalization medium (EBM-2) at 37°C for 1h. Finally, HUVEC were incubated in acid wash (0.5M NaCl/0.2M acetic acid) for 4 min at 4°C to remove remaining labeled VE-cadherin on the cell surface, then washed with PBS+ and fixed. For quantification, internalized VE-cadherin area was measured in Fiji, then normalized to total cell area for at least 9 cells per experiment.

### Microtubule analysis

For nuclear microtubule analysis, an ROI was drawn in Fiji over the nucleus. Fiji was used to threshold α-tubulin signal, and the area of the signal was measured within the ROI. This was performed on 8 cells per field of view, with 3-6 fields per condition per experiment. For high-resolution microtubule analysis, high-resolution confocal images were acquired with a Zeiss 880 Confocal with AiryScan FAST microscope with GaAsP detector and camera (Zeiss) using a Plan-Apo 63x oil immersion objective (NA 1.40) and 488nm and 561nm lasers. Imaging was performed at RT, and images were acquired with the Zeiss 880 software and with the AiryScan detector. Images were then processed with AiryScan. All image analysis, including Z-stack compression, was performed in Fiji. For microtubule density, an ROI was drawn at the MTOC and the cell periphery. Fiji was used to threshold α-tubulin signal, and the area of the signal was measured within the ROI. This was performed on at least 10 cells per condition per experiment. For junction analysis, the number of contacts between α-tubulin and VE-cadherin were counted and normalized to the junction length. At least 15 junctions were measured per condition per experiment.

### Microtubule tip tracking

HUVEC were infected with an EB3-mCherry Lentivirus (Kushner et al., 2014) for 24h to visualize microtubule comets. Following infection, HUVEC were incubated with siRNAs for NT or SUN1. For live imaging, cells were incubated at 37°C in a stage-top incubator (TOKAI HIT, WSKM). Images were acquired on an Andor XD spinning disk confocal microscope based on a CSU-X1 Yokogawa head with an Andor iXon 897 EM-CCD camera. A 561nm laser and FF01-607/36 emission filter were used. Images were acquired with a 470ms exposure and 32ms readout time for 2 min (240 frames) using a UPlanSApo 60x silicone-oil immersion objective (NA 1.30) and Metamorph software. Microtubules at the cell periphery were tracked using the Manual Tracking plugin in Fiji and were tracked for at least 60 frames (30 sec). Track information was acquired from the x and y coordinates using a custom algorithm in Visual Basic in Excel provided by Dan Buster at the University of Arizona. 10 microtubule tracks were measured per cell.

### GEF-H1 analysis

For GEF-H1 localization analysis, an ROI was drawn at the periphery of the cell, and the α-tubulin signal was used to create a mask. Mean signal intensity was then measured for GEF-H1 within the α-tubulin mask and outside of it, and a ratio was taken. At least 9 cells were analyzed per condition per experiment.

### Statistics

Student’s two-tailed unpaired *t-*test was used to determine statistical significance in experiments with 2 groups and one-way ANOVA with Tukey’s multiple comparisons test was used in experiments with 3 groups. For thrombin, blebbistatin, Y-27632, and nocodazole experiments, two-way ANOVA with Tukey’s multiple comparisons test was used to determine statistical significance. Χ^2^ was used for categorical data. For box and whisker plots, boxes represent the upper quartile, lower quartile, and median; whiskers represent the minimum and maximum values. Statistical tests and graphs were made using the Prism 9 software (GraphPad Software).

### Data availability

The RNA-seq data that support the findings of this study are available in the Gene Expression Omnibus (GEO) under the accession number GSE213099.

## ACKNOWLEGMENTS

We thank Michelle Altemara and staff at the Zebrafish Aquaculture Core and Kaitlyn Quigley for fish room support, Aaron Friedman and Caroline Crater for mouse room support, and Yosuke Mukouyama, Nick Buglak, and Bautch Lab members for critical discussion and feedback. We also thank Dan Buster for the software used in microtubule tip tracking experiments and Angelika Noegel for sharing RNASeq data with us. Airy Scan imaging was performed at the UNC Hooker Imaging Core Facility and microtubule tip tracking imaging was performed with Pablo Ariel at the UNC Microscopy Services Laboratory, supported in part by P30 CA016086 Cancer Center Core Support Grant to the UNC Lineberger Comprehensive Cancer Center.

## AUTHOR CONTRIBUTIONS

Danielle B Buglak (DBB) and Victoria L Bautch (VLB) conceptualized the work; DBB, Ariel L Gold (ALG), Allison P Marvin (APM), Shea N Ricketts (SNR), Molly R Kulikauskas (MRK), Andrew Burciu (AB), Karina Kinghorn (KK), Morgan Oatley (MO), Natalie T Tanke (NTT), Pauline Bougaran (PB), and Bryan N Johnson (BNJ) performed and analyzed experiments; Ziqing Liu (ZL) analyzed RNASeq data; DBB and VLB wrote and edited the manuscript; Celia E Shiau (CES) provided oversight on zebrafish experiments; Stephen L Rogers (SLR) provided oversight and discussion on microtubule and GEF-H1 experiments; VLB provided study supervision and oversight.

