## Supplementary Material for "Nuclear SUN1 Stabilizes Endothelial Cell Junctions via Microtubules to Regulate Blood Vessel Formation"

Buglak et al.

### SUPPLEMENTARY MATERIAL

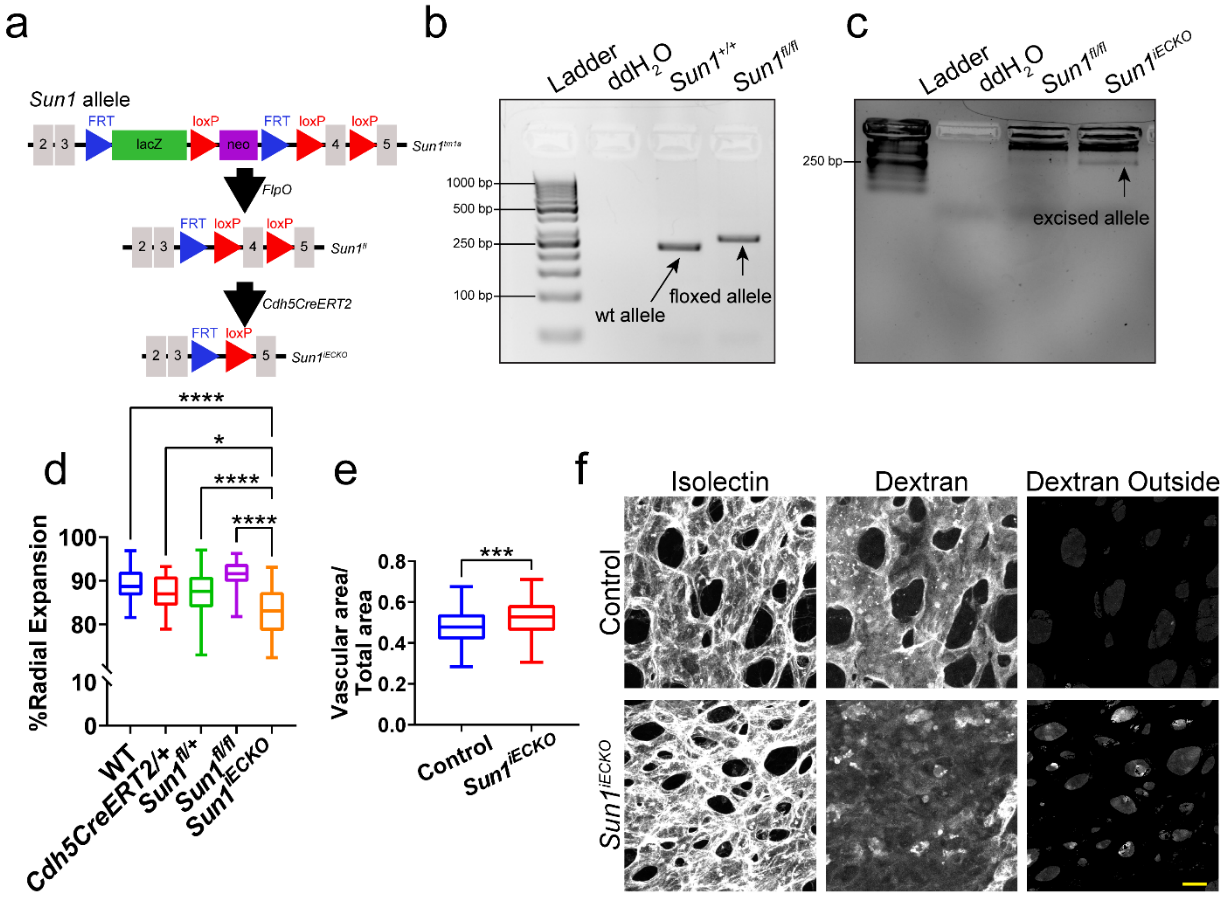


**Figure 1-figure supplement 1. Loss of *Sun1* in the postnatal retina leads to altered sprouting and barrier function.**
**(a)** Schematic showing strategy for generation of the *Sun1* floxed allele and subsequent Cre-mediated excision of exon 4 of the *Sun1* allele. **(b)** Agarose ethidium bromide gel showing PCR bands specific for WT or $Sun1^{fl}$ allele. **(c)** Agarose ethidium bromide gel showing PCR band specific for the excised *Sun1* allele from mouse lung tissue. **(d)** Graph of radial expansion from **Figure 1c** with genotypes broken out. *, $p<0.05$; ****, $p<0.0001$ by one-way ANOVA with Tukey's multiple comparisons test. **(e)** Graph of vascular density of combined ROIs from **Figure 1D**. n=171 ROIs from 27 retinas (controls) and 75 ROIs from 12 retinas ($Sun1^{iECKO}$) from 3 independent litters. ***, $p<0.001$ by student's two-tailed unpaired *t*-test. **(f)** Representative images showing dextran labeling within P7 mouse retinas with indicated genotypes. Scale bar, 20μm.

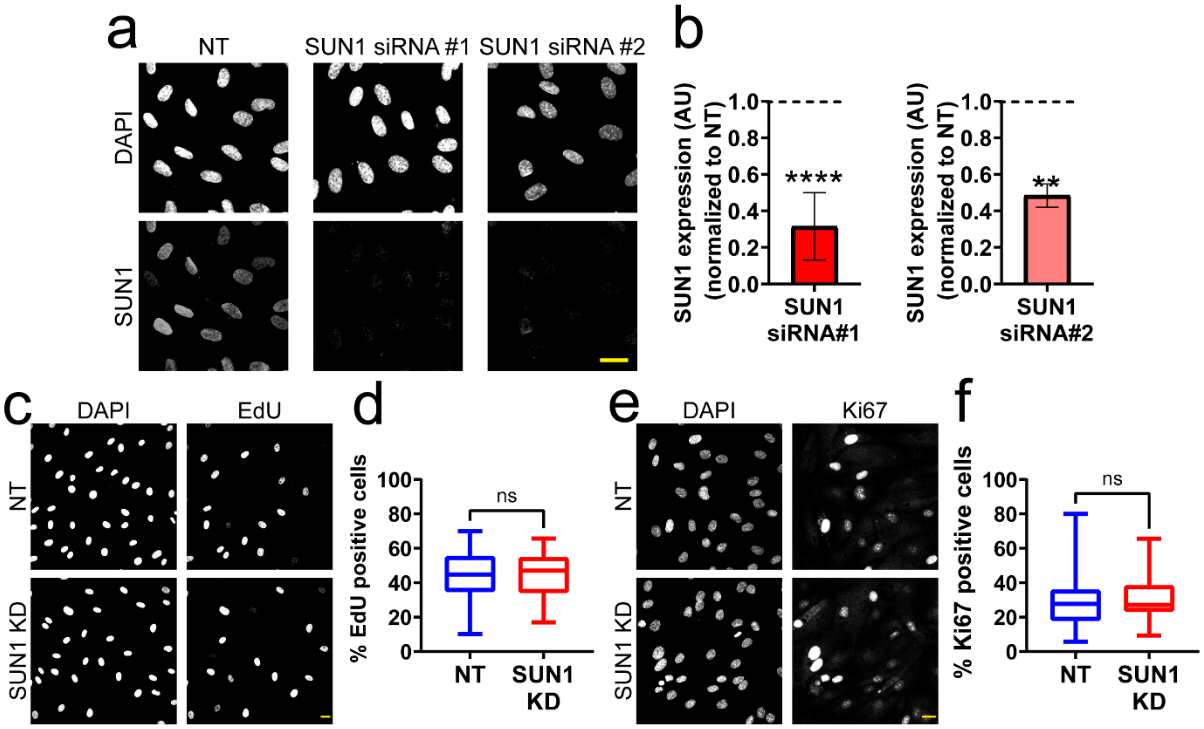


**Figure 2-figure supplement 1. SUN1 is nuclear localized in endothelial cells and does not regulate proliferation.**
**(a)** Representative images of HUVEC with indicated siRNAs and stained with the indicated antibodies. Endothelial cells were stained for DAPI (DNA) and SUN1. Scale bar, 20µm. **(b)** Quantification of SUN1 expression shown in **a**. Expression is normalized to NT. n=9 replicates (SUN1 siRNA#1) and 2 replicates (SUN1 siRNA#2). **, $p<0.01$; ****, $p<0.0001$ by student's two tailed unpaired *t*-test. **(c)** Representative images of HUVEC with indicated siRNAs and EdU incorporation. Endothelial cells were stained for DAPI (DNA) and EdU. Scale bar, 20µm. **(d)** Quantification of percent EdU positive cells shown in **c**. n=30 ROIs (NT) and 30 ROIs (SUN1 KD) compiled from 2 replicates. ns, not significant by student's two-tailed unpaired *t*-test. **(e)** Representative images of HUVEC with indicated siRNAs and Ki67 staining. Endothelial cells were stained for DAPI (DNA) and Ki67. Scale bar, 20µm. **(f)** Quantification of percent Ki67 positive cells shown in **e**. n=24 ROIs (NT) and 15 ROIs (SUN1 KD) compiled from 3 replicates. ns, not significant by student's two-tailed unpaired *t*-test.

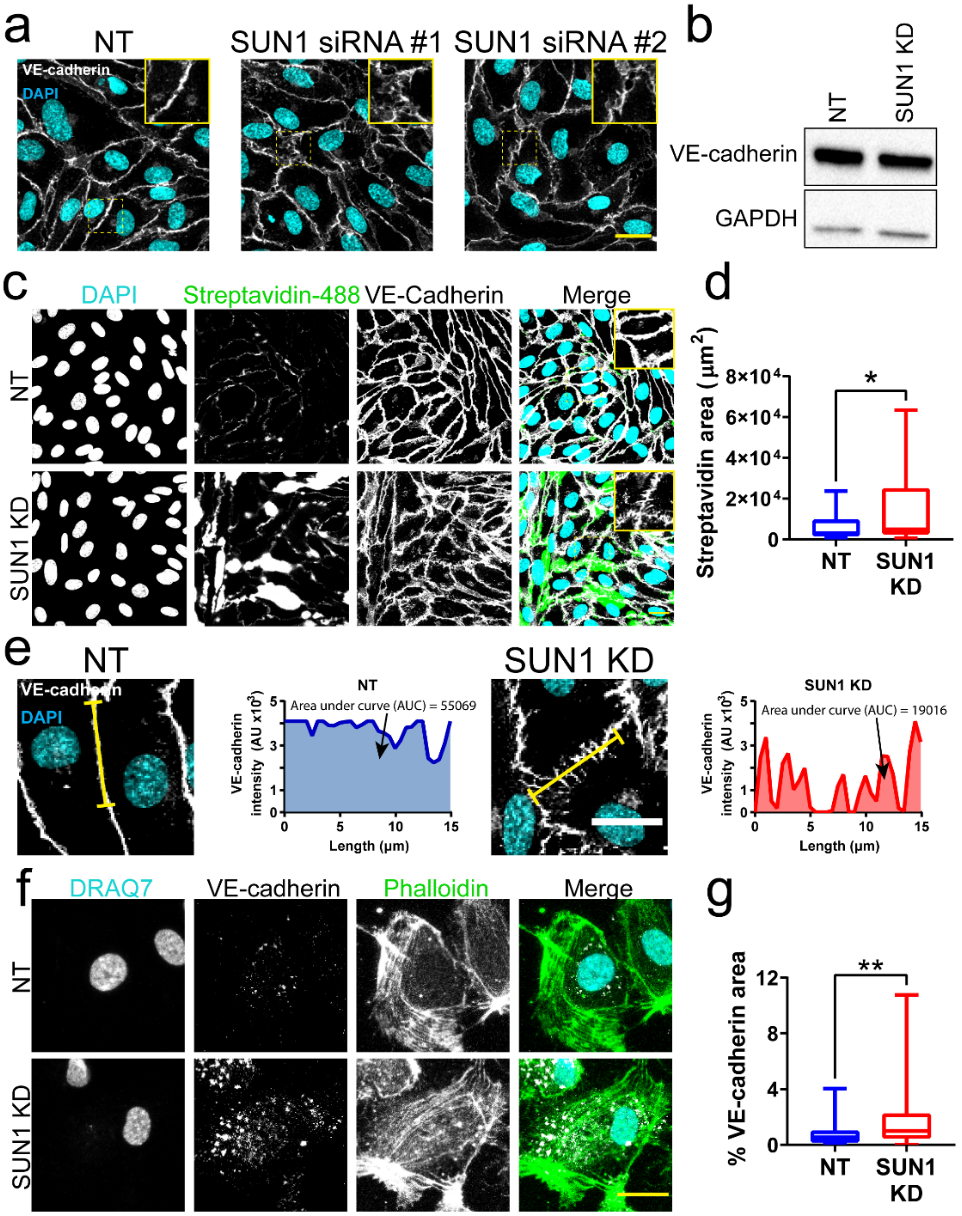


**Figure 4-figure supplement 1. SUN1 regulates endothelial cell barrier function.**
**(a)** Representative images of HUVEC with indicated siRNAs. Endothelial cells were stained for DAPI (cyan, DNA) and VE-cadherin (white, junctions). Insets show junctions. Scale bar, 20μm. **(b)** Representative western blot showing levels of VE-cadherin protein in HUVEC with indicated siRNAs. GAPDH was used as a loading control. **(c)** Representative images of HUVEC with indicated siRNAs cultured on biotinylated fibronectin and treated with streptavidin upon confluence. Endothelial cells were stained for DAPI (cyan, DNA), Streptavidin (green), and VE-cadherin (white, junctions). Insets show junctions. Scale bar, 20μm. **(d)** Quantification of streptavidin area shown in **c**. n=29 ROIs (NT) and 29 ROIs (SUN1 KD) compiled from 4 replicates. *, $p<0.05$ by student's two-tailed unpaired *t*-test. **(e)** Representative images and graphs of HUVEC with indicated siRNAs showing VE-cadherin line scan quantification. Endothelial cells were stained with DAPI (cyan, DNA) and VE-cadherin (white, junctions). Yellow line indicates where line scan was taken. Arrows point to the area under the curve (AUC). Scale bar, 20μm. **(f)** Representative images of HUVEC with indicated siRNAs showing VE-cadherin staining after internalization assay. Endothelial cells were stained for DRAQ7 (cyan, DNA), VE-cadherin (white, junctions), and Phalloidin (green, actin). Scale bar, 20μm. **(g)** Quantification of area of internalized VE-cadherin in **f.** n=57 cells (NT) and 82 cells (SUN1 KD) compiled from 3 replicates. **, $p<0.01$ by student's two-tailed unpaired *t*-test.

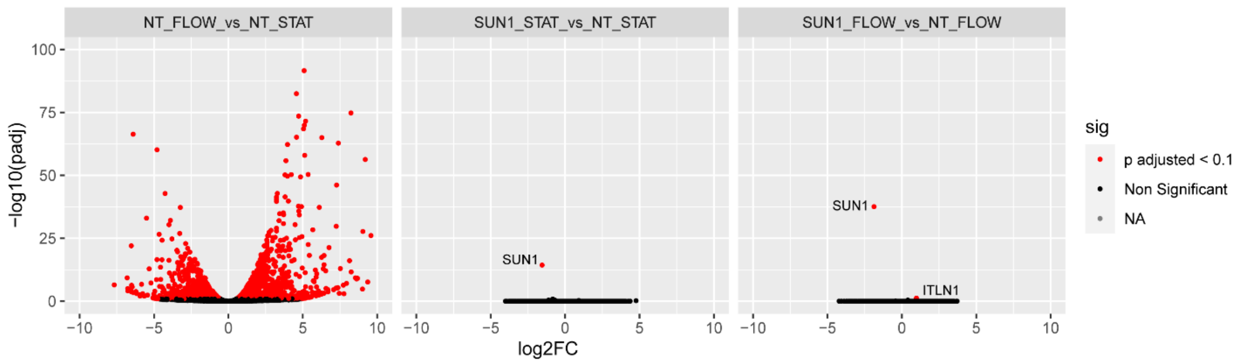


**Figure 5-figure supplement 1. SUN1 does not affect transcription in endothelial cells.** Volcano plots showing transcriptional changes from bulk RNASeq data in NT (non-targeting) under flow vs NT under static conditions (left), SUN1 KD under static vs NT under static conditions (middle), and SUN1 KD under flow vs NT under flow conditions (right). Differentially expressed genes were determined by DESeq2 and considered significant with an adjusted p-value <0.01. Red dots, significant genes; black dots, non-significant genes; green dots, NA.

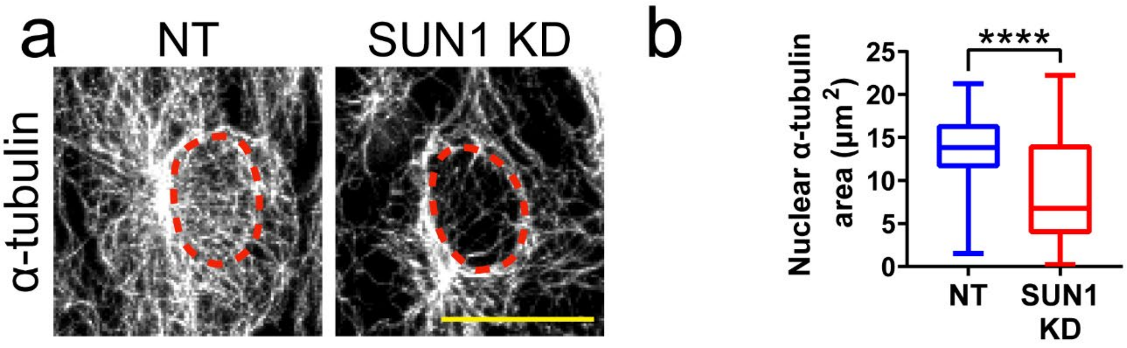


**Figure 5-figure supplement 2. SUN1 regulates the microtubule cytoskeleton in endothelial cells.**
**(a)** Representative images of HUVEC with indicated siRNAs showing changes in microtubules at nucleus. Endothelial cells were stained for α-tubulin (white, microtubules). Red dashed line shows outline of nucleus. Scale bar, 20μm. **(b)** Quantification of nuclear α-tubulin area shown in **a**. n=80 cells (NT) and 72 cells (SUN1 KD) compiled from 3 replicates. ****, $p$<0.0001 by student's two-tailed unpaired $t$-test.

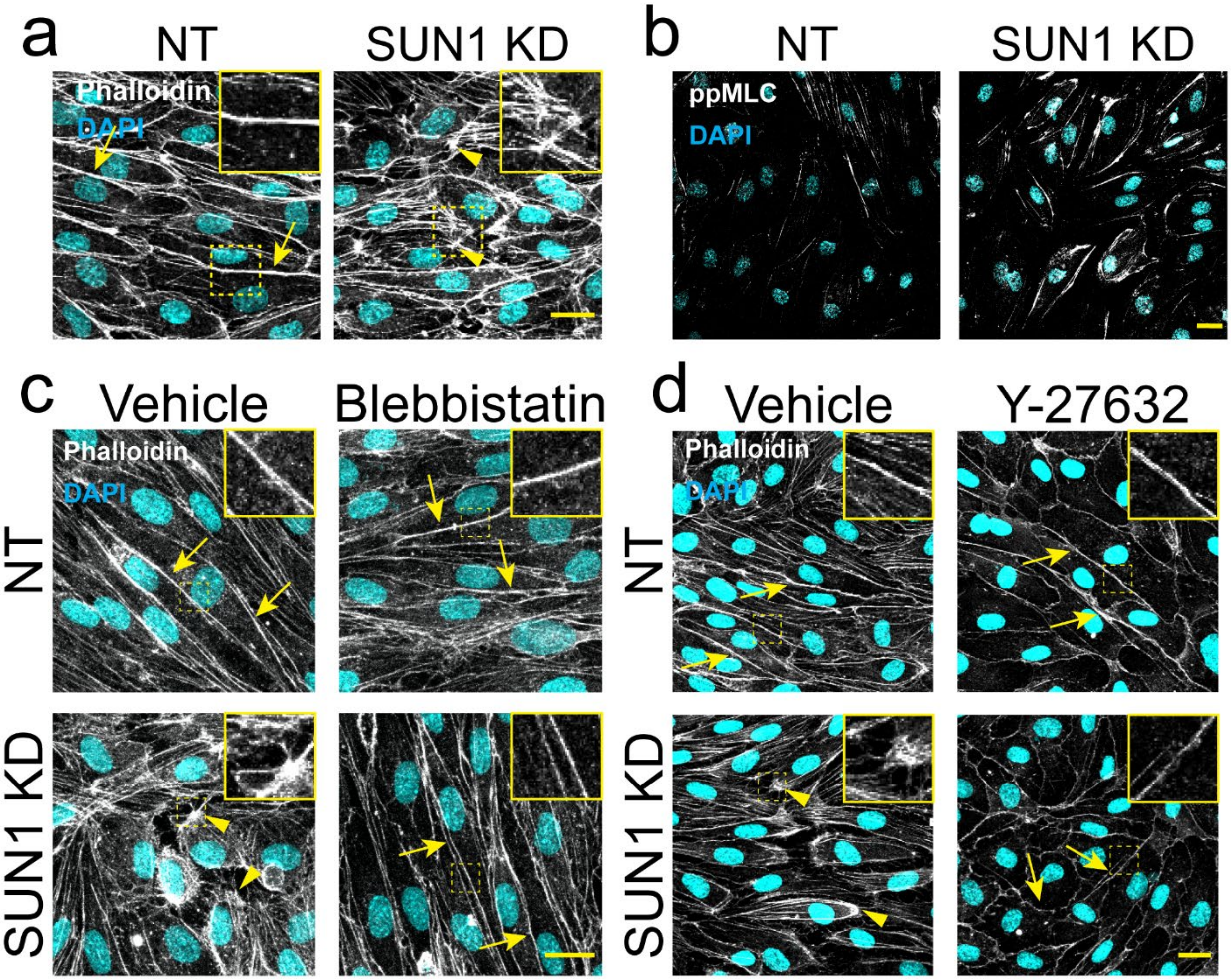


**Figure 6-figure supplement 1. SUN1 regulates the actin cytoskeleton in endothelial cells.** **(a)** Representative images of HUVEC with indicated siRNAs showing changes in actin structures at the cell periphery. Endothelial cells were stained for DAPI (cyan, DNA) and VE-cadherin (white, junctions). Insets show actin structures at periphery. Scale bar, 20μm. **(b)** Representative images of HUVEC with indicated siRNAs stained for ppMLC. Endothelial cells were stained for DAPI (cyan, DNA) and ppMLC (white). Scale bar, 20μm. **(c)** Representative images of HUVEC with indicated siRNAs and indicated treatments. Endothelial cells were stained for DAPI (cyan, DNA) and Phalloidin (white, actin). Insets show actin structures at periphery. Scale bar, 20μm. **(d)** Representative images of HUVEC with indicated siRNAs and indicated treatments. Endothelial cells were stained for DAPI (cyan, DNA) and Phalloidin (white, actin). Insets show actin structures at periphery. Scale bar, 20μm. Arrows denote cortical actin; arrowheads denote radial actin bundles.

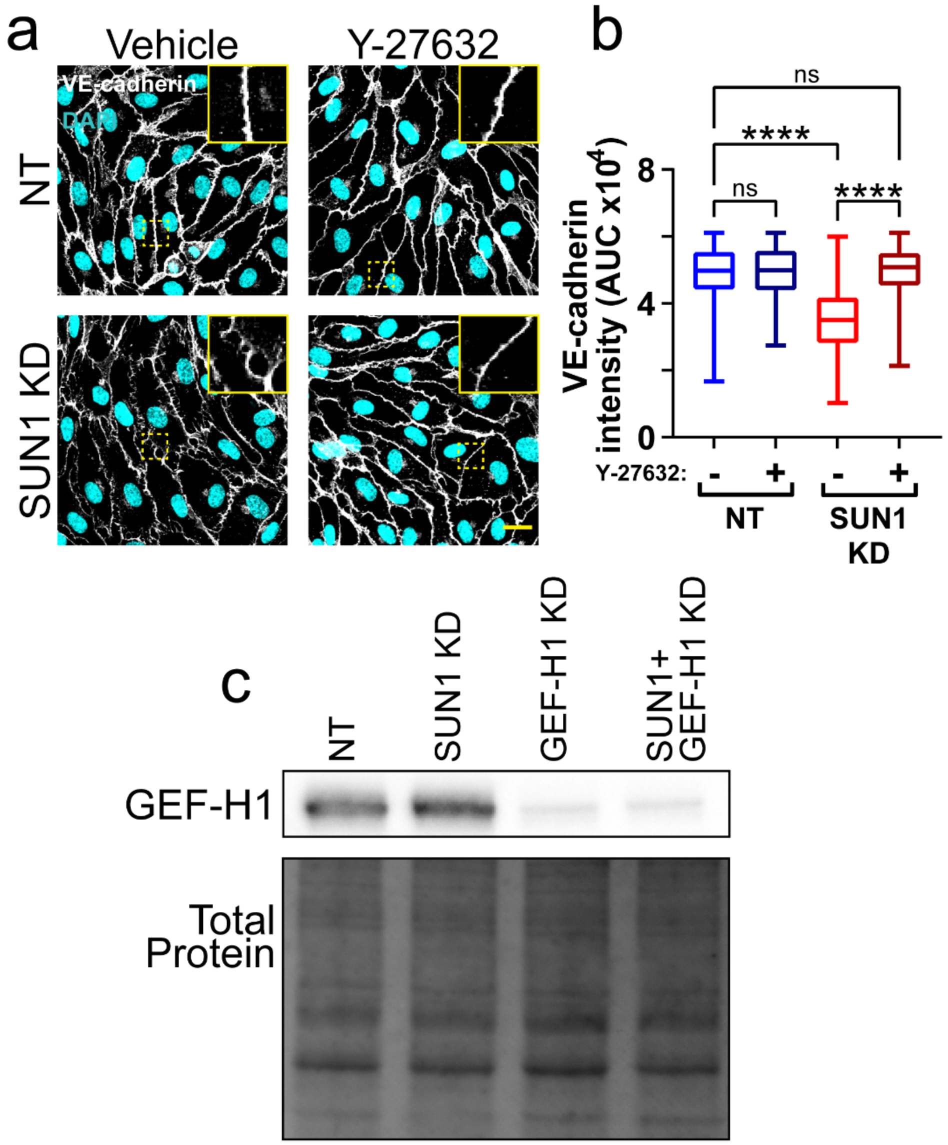


**Figure 6-figure supplement 2. SUN1 acts through Rho signaling and GEF-H1 to affect junctions.**
**(a)** Representative images of HUVEC with indicated siRNAs. Endothelial cells were stained for DAPI (cyan, DNA) and VE-cadherin (white, junctions). Insets show junctions. Scale bar, 20µm. **(b)** Quantification of VE-cadherin line scans for treatments shown in **a**. n=185 junctions (NT, vehicle), 165 junctions (NT, Y-27632), 175 junctions (SUN1 KD, vehicle), and 173 junctions (SUN1 KD, Y-27632) compiled from 3 replicates. Ns, not significant; ****, *p*<0.0001 by two-way ANOVA with Tukey's multiple comparisons test. **(c)** Representative western blot showing total levels of GEF-H1 protein in HUVEC with indicated siRNAs. Total protein was used as a loading control.

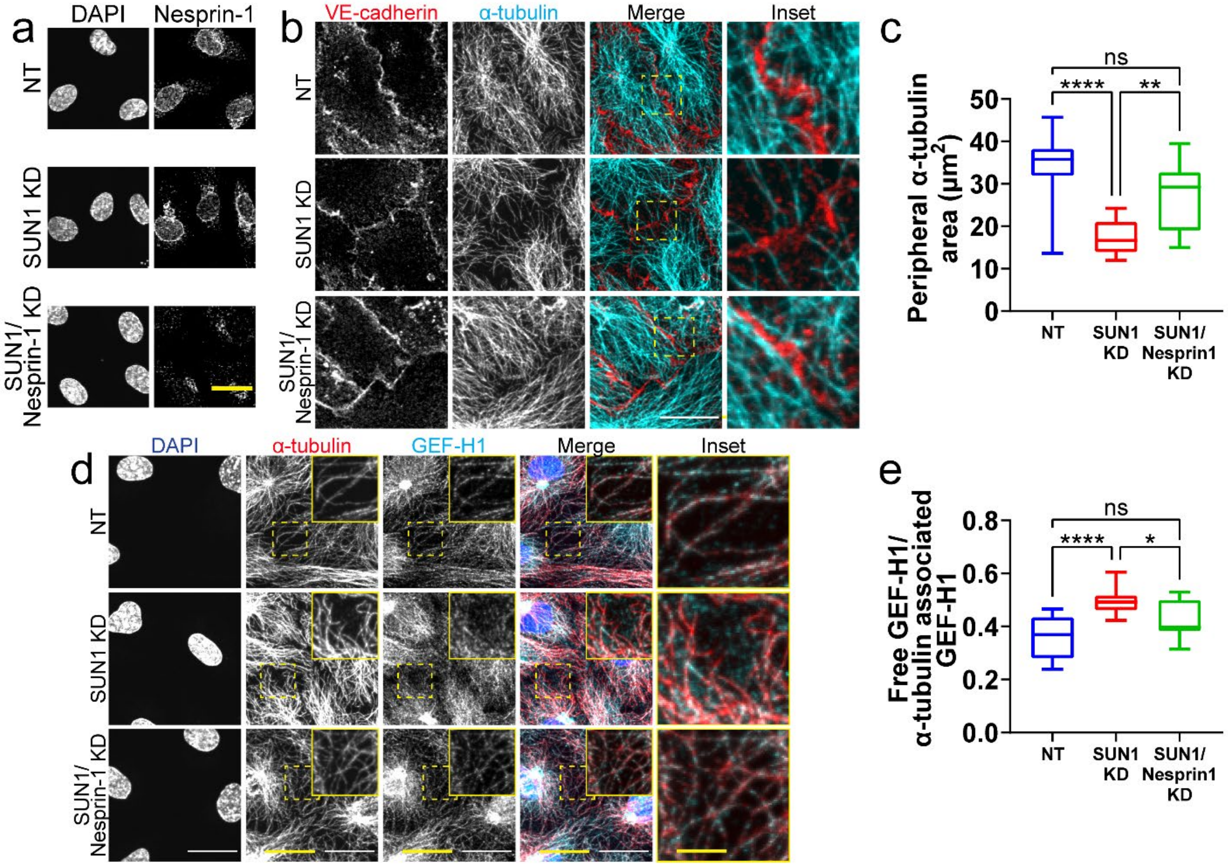


**Figure 7-figure supplement 1. SUN1 exerts its effects on junctions through nesprin-1.** **(a)** Representative images of HUVEC with indicated siRNAs and indicated antibodies. Endothelial cells were stained for DAPI (DNA) and nesprin-1. Scale bar, 20μm. **(b)** Representative images of HUVEC with indicated siRNAs. Endothelial cells were stained for α-tubulin (cyan, microtubules) and VE-cadherin (red, junctions). Insets show α-tubulin contacts at junctions. Scale bar, 20μm. **(c)** Quantification of peripheral α-tubulin area shown in **b**. n=12 cells (NT), 12 cells (SUN1 KD), and 12 cells (SUN1/Nesprin-1 KD) from 1 representative replicate. ns, not significant; **, $p$<0.01; ****, $p$<0.0001 by one-way ANOVA with Tukey's multiple comparisons test. **(d)** Representative images of HUVEC with indicated siRNAs. Endothelial cells were stained for DAPI (blue, DNA), α-tubulin (red, microtubules), and GEF-H1 (cyan). Insets show α-tubulin and GEF-H1 colocalization. Scale bar, 20μm. **(e)** Quantification of free GEF-H1 normalized to α-tubulin associated GEF-H1 shown in **d**. n=12 cells (NT), 12 cells (SUN1 KD), and 12 cells (SUN1/Nesprin-1 KD) from 1 representative replicate. ns, not significant; *, $p$<0.05; ****, $p$<0.0001 by one-way ANOVA with Tukey's multiple comparisons test.

**Movie 1. Control endothelial cells elongate in 3D sprouting assay.**

3D sprouting angiogenesis of control (NT) HUVEC over 60h, showing elongation of NT sprouts. Scale bar, 50µm. Frames acquired every 30 min.

**Movie 2. SUN1-depleted endothelial cells retract in 3D sprouting assay.**

3D sprouting angiogenesis of SUN1 KD sprouts over 60h, showing retraction of SUN1 KD sprouts. Scale bar, 50µm. Frames acquired every 30 min.

**Movie 3. Control zebrafish have normal ISV growth.**

Movie taken from 26-36 hpf in *Tg(fli:LifeAct-GFP)* zebrafish embryos injected with a NT morpholino, showing elongation of ISVs and connection to the DLAV. A, anterior; P, posterior. Scale bar, 20µm. Frames acquired every 15 min.

**Movie 4. Loss of SUN1 in zebrafish leads to abnormal ISV growth.**

Movie taken from 26-36 hpf in *Tg(fli:LifeAct-GFP)* zebrafish embryos injected with *a sun1b* morpholino, showing an ISV that fails to elongate and connect to the DLAV and an ISV that elongates but does not connect to the DLAV. A, anterior; P, posterior. Arrow points to ISV that does not elongate. Scale bar, 20µm. Frames acquired every 15 min.

**Movie 5. Control endothelial cells have normal microtubule dynamics.**

Movie taken for 120 sec in HUVEC with NT siRNA labeled with EB3-mCherry. Quantified microtubule tracks are indicated. Scale bar, 5µm. Frames acquired every 500 ms.

**Movie 6. Loss of SUN1 in endothelial cells leads to impaired microtubule dynamics.**

Movie taken for 120 sec in HUVEC with SUN1 siRNA labeled with EB3-mCherry. Quantified microtubule tracks are indicated. Scale bar, 5µm. Frames acquired every 500 ms.
